# Modelling interpretable patient-level representations from structured and simple multimodal data

**DOI:** 10.64898/2026.08.31.748244

**Authors:** Kazimierz Oksza-Orzechowski, Małgorzata Łazęcka, Łukasz Koperski, Damian Wójtowicz, Marcin Możejko, Daniel Schulz, Robin Liechti, Flavia Marzetta, Marie Morfouace, Henoch S. Hong, Stéphanie Tissot, Bernd Bodenmiller, Eike Staub, Ewa Szczurek

**Affiliations:** Faculty of Mathematics, Informatics and Mechanics, University of Warsaw, Warsaw, Poland; Department of Pathology, Medical University of Warsaw, Warsaw, Poland; Centre for Credible AI, Warsaw University of Technology, Warsaw, Poland; Department of Quantitative Biomedicine, University of Zurich, Zurich, Switzerland; Institute of Molecular Health Sciences, ETH Zurich, Zurich, Switzerland; Vital-IT, SIB Swiss Institute of Bioinformatics, Lausanne, Switzerland; EORTC HQ, Brussels, Belgique; Gustave Roussy, Villejuif, France; The healthcare organization of Merck KGaA, Darmstadt, Germany; Centre hospitalier Universitaire Vaudois, Lausanne, Switzerland; Institute of AI for Health, Helmholtz Munich, Neuherberg, Germany

## Abstract

Patient cohort profiling increasingly includes structured views for multiple modalities, such as single-cell RNA sequencing, spatial transcriptomics or proteomics, and histology, each providing multiple subobservations per patient, including single cells, spatial spots or patches. To model such data along with simple patient-level views, current multimodal integration methods typically rely on separately precomputed summaries and fail to fully leverage information in structured views. Here we present FACTMx, a variational framework that jointly models structured and simple views to learn interpretable patient-level representations. FACTMx couples latent patient factors with subobservation clustering and per-patient component proportions, enabling direct interpretation and downstream association analyses. The framework supports different structured-view mixture assumptions, including topic- and Gaussian-structured data, while retaining modular encoder-decoder parameterisations. In simulations spanning sparse and dense dependencies and multiple noise regimes, FACTMx improved reconstruction, integration and recovery of structured components relative to previous methods. Applied to non-small cell lung cancer cohorts, FACTMx captured survival-associated latent signals linked to immune microenvironments, gene expression pathways and spatially coherent histological patterns. In a longitudinal coronary syndrome cohort, FACTMx highlighted an outcome-associated axis connected to ejection-fraction change, immune cell states, soluble mediators and cardiac injury markers. These results support joint structured-simple modelling for interpretable multimodal patient stratification.

## Main

Biological processes are visible across multiple levels of organisation. Immune response, for example, can be detected in gene expression programs at single-cell resolution, in spatially organised microenvironments within tissue, and in bulk molecular profiles or clinical variables measured at the patient level [1–4]. Consequently, in-depth study of such processes requires methods that combine signals from all available levels into patient-level representations while preserving interpretability. This need has become more pressing with the rapid growth of multimodal cohorts and profiling technologies, which routinely generate *multi-level* data, with combinations of *structured views* with multiple *subobservations* per patient, as well as *simple views* carrying a single vector of values per patient [5–8]. Structured view subobservations can be, for example, cells from single-cell RNA sequencing (scRNA-seq), spots from spatial transcriptomics, or patches from Hematoxylin and Eosin (H&E) histology slides. Simple view vectors can be blood test values or bulk RNA-seq measurements per patient.

Despite substantial progress in multimodal learning, methods that directly support this type of integrative multilevel data modelling remain limited. Supervised multimodal approaches based on multiple instance learning (MIL) or deep fusion can aggregate lower-level observations and combine them with additional modalities, but are typically optimised for one predefined prediction target, such as diagnosis, prognosis, or treatment response, rather than learning a general-purpose patient representation for downstream exploration [9, 10]. In practice, patient-level representations are therefore often obtained using a two-step procedure. In the first step, structured modalities are summarised independently of each other, for example, by finding clusters of subobservations using latent Dirichlet allocation (LDA), Gaussian mixture models or graph-based clustering such as Leiden, and creating per-modality *patient summaries* corresponding to cluster abundances [11, 12]. For example, single cells can be clustered into celltypes based on their scRNA-seq profiles, and a patient can be summarised as a vector of celltype frequencies. In the second step, patient summaries are treated as further simple modalities and integrated with remaining simple modalities using an autoencoder or a factor model such as MOFA [13, 14]. Although effective in some settings [5, 6, 15–17], such pipelines decouple the construction of structured summaries from multimodal integration, which limits the extent to which information can be shared across levels and modalities. An important step towards joint modelling of structured and simple modalities is FACTM [18], which combines factor analysis with correlated topic modelling and infers both components simultaneously. This coupling improves integration relative to sequential procedures and yields interpretable latent factors. However, FACTM is limited to topic-model structure and linear cross-modal dependencies, which disqualifies it in settings requiring other mixture assumptions for structured modalities or nonlinear dependencies. Furthermore, its explicit graph formulation hinders extensibility, since adding support for new distributions requires deriving and recalculating the posterior update equations.

In this work, we introduce FACTMx, a general framework for multimodal representation learning based on flexible variational autoencoding of both simple and structured data. In the model, the patient-level representations are inferred jointly with the parameters behind structured view mixtures, improving both the quality of multimodal integration and the correspondence between patient-level factors and structured observations. FACTMx allows structured components to be instantiated with different mixture models and supports nonlinear dependencies through modular encoder–decoder parameterisations, while retaining patient-level interpretability that facilitates downstream association analyses. We evaluate FACTMx on simulated and experimental datasets and show that it improves modality integration and downstream interpretability over existing approaches.

## Results

### FACTMx model

#### Overview

FACTMx takes as input multimodal measurements of a cohort and integrates them into a shared, interpretable representation per sample. We distinguish two types of input data based on their structure (Fig. 1**a**). *Simple views* are modalities for which each sample is described by a single feature vector; for example, bulk RNA-seq counts, mutational signature exposures, or clinical variables. *Structured views* are modalities for which each sample is described by a collection of feature vectors, hereafter called *subobservations*; for example, single cells profiled from one patient, image patches extracted from one histology slide, or cell niches segmented from one tissue section. A sample can be measured across any combination of simple and structured modalities, and the number of subobservations per sample is allowed to vary. From these inputs, FACTMx jointly infers three coupled outputs that together describe a sample. The first is a low-dimensional *patient-level representation*: a vector summarising the sample across all modalities. We refer to its dimensions as *factors*, and each factor is interpretable in the sense that its effect on every input modality can be read off directly from the model’s weights (see below). The second output, defined for each structured modality, is a clustering structure with a fixed number of components. These components describe, for example, niche types in spatial imaging or cell states in single-cell data and are interpretable by analysing either their parameters in the model or the subobservations assigned to them. The third output, derived from this clustering, is the set of per-sample component proportions: for each structured modality, the fraction of that sample’s subobservations assigned to each component. Together, these outputs serve as a multimodal embedding of the cohort that can be used directly in downstream analyses such as survival modelling, patient stratification, or identification of clinical and molecular associations.

**Figure 1:**
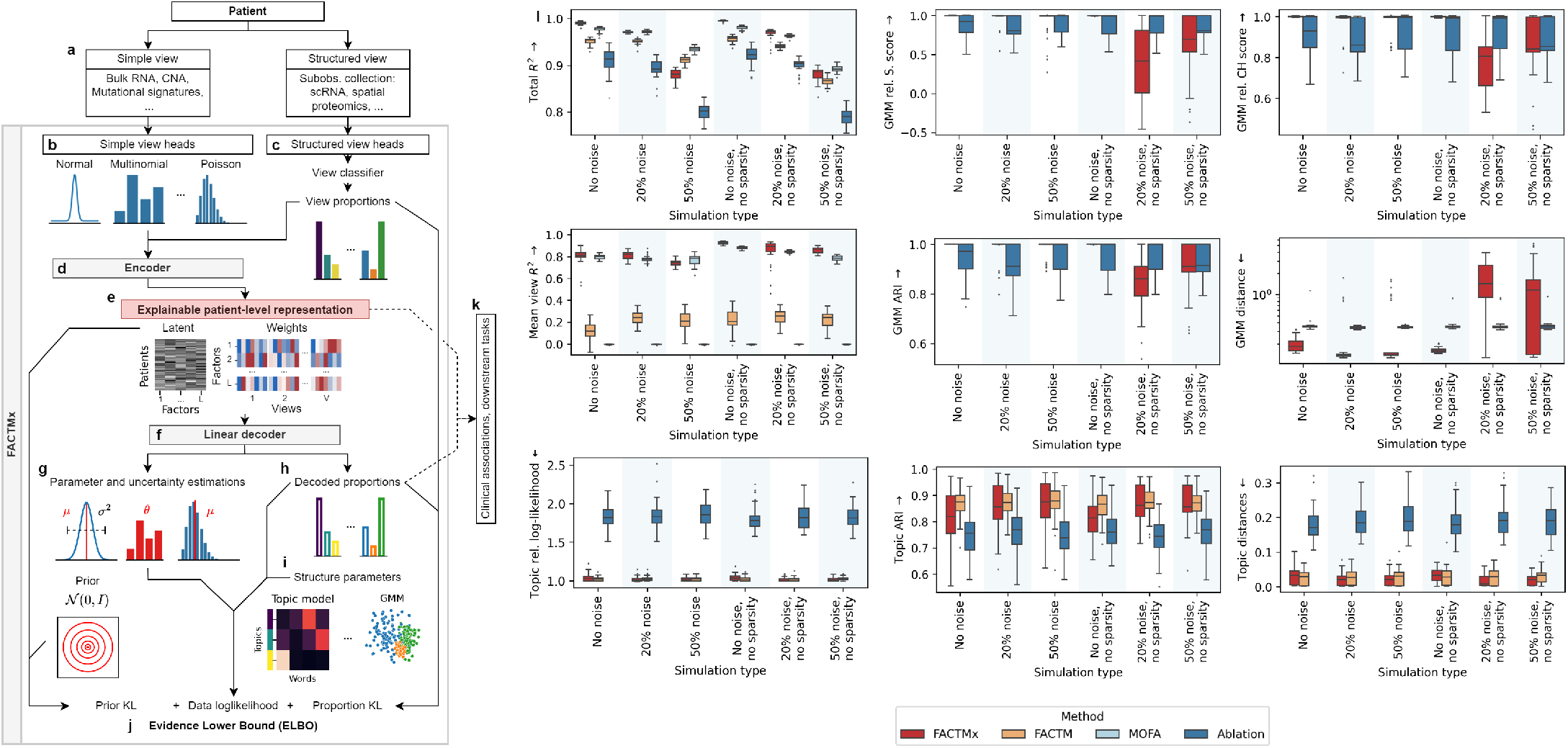
FACTMx pipeline overview **a-k**: **a** an observation point (e.g. a patient) is measured across a range of modalities, either simple or structured; each modality is handled with an appropriate head, either simple (**b**) or structured (**c**); **d** an encoder takes as input transformed data from **b** and structured patient summaries from initial classifiers in **c**; **e-f** the latent space of the model is designed to provide interpretable patient-level representations through the use of a linear decoder, which decodes distribution parameters for simple views (**g**) and patient summaries for structured views (**h**); **i** each structured head contains trainable parameters of its components; **j** the model is trained by optimising ELBO, which consists of a prior KL divergence term between the patient representatations in **e** and an uninformative prior, data log-likelihood obtained by using parameters from **g** and combining the summaries and parameters from **h-i**, and a summary KL term between decoded summaries **d** and initial summaries from **c**; **k** we use patient representation (**e**) and decoded patient summaries (**h**) to improve downstream analysis tasks. **l** Comparison of FACTMx with competitor models on simulated datasets; methods are grouped under ablation for metrics where their application process is identical. *Subobs* – subobservations, *KL* – Kullback-Leibler, *rel*. – relative, *S*. – silhouette, *CH* – Calinski-Harabasz.

FACTMx (Fig. 1**b-j**, Methods) is a variational autoencoder with one module, named a *head*, per input modality (Fig. 1**a**). A *simple head* models its modality with a parametric distribution (e.g. Normal, Multinomial, Poisson) whose parameters are decoded from the patient representation, optionally combined with directly optimised parameters (Fig. 1**b**). A *structured head* models its modality with a mixture (e.g. a topic model or a mixture of Gaussians) and additionally trains a classifier that assigns each subobservation to a mixture component (Fig. 1**c**). Averaging these assignments within a sample yields a primary estimate of its patient component proportions. The encoder (Fig. 1**d**) takes the simple-data inputs and the patient component proportions from structured views and outputs the parameters of a variational posterior over the patient representation (Fig. 1**e**). A linear decoder (Fig. 1**f**) maps this representation back to the parameters of each simple view’s data distribution (Fig. 1**g**) and to a decoded estimate of each patient component proportions (Fig. 1**h**). Because the mapping is linear, the contribution of every factor to each modality is given directly by a row of the decoder matrix, which is what makes factors interpretable. Finally, the decoded patient component proportions are combined with trainable per-component parameters of each structured head (Fig. 1**i**), for example, topic word profiles or Gaussian means and covariances, to define subobservation likelihoods.

The model is trained end-to-end by variational inference (Fig. 1**j**), comprising three terms: Kullback–Leibler (KL) divergence between the variational posterior over the patient representation and a standard normal prior and the data log-likelihood across all simple and structured views (both typical for variational autoencoders), as well as a novel loss term, namely a KL divergence between the component proportions derived bottom-up from the structured-view classifiers and the component proportions decoded top-down from the patient representation. As a result, the clustering of subobservations within each structured view, the patient component proportions, and the patient representation are inferred jointly rather than in sequence. This is one of the key features distinguishing FACTMx from two-step pipelines that cluster each structured modality before integration. After training, patient representations, factor loadings, subobservation cluster assignments, and patient component proportions are extracted from the model and used directly in downstream analyses (Fig. 1**k**).

#### Synthetic benchmark

We benchmarked FACTMx on simulated data against three competitors: a vanilla autoencoder (AE; ablation), MOFA, and FACTM. Each simulated dataset comprised both simple and structured modalities, the latter including two topic-style views and one mixture-of-Gaussians view. Since neither AE nor MOFA can ingest collections of subobservations directly, and FACTM does not support mixture-of-Gaussians views, we first summarised every structured modality into a per-patient component-proportion vector using a 10-component LDA for topic modalities and a 10-component Gaussian mixture model (GMM) for the mixture-of-Gaussians modality, and used these summaries as their structured-view inputs (see Methods). AE additionally concatenated all simple-view and summarised-structured-view inputs into a single vector and was trained with a linear encoder-decoder and an RMSE loss. We considered six simulation scenarios (see Methods) defined by two characteristics: dependency sparsity (sparse or not) and noise level (0%, 20%, 50%). Briefly, sparse dependency means each latent direction influences only a subset of the patient’s data views, and noise level is the fraction of variance in patient simple views not explained by their latent representations. We evaluate the models on nine metrics: total data reconstruction (total *R*^2^); data integration (mean *R*^2^ across views); silhouette (S.) and Calinski–Harabasz (CH) scores of the reconstructed GMM relative to those of the generative GMM; adjusted Rand index (ARI) between the reconstructed and original clusterings in the GMM and topic views; distances between the reconstructed and generative component parameters of the GMM and topic views; and the log-likelihood of subobservations in the topic views relative to that of the generative model.

Across 30 datasets per scenario (Fig. 1**l**), FACTMx is the best or tied-best method on every metric in the majority of scenarios, with the gains taking a different form against each competitor. Against the AE ablation, FACTMx improves nearly every metric and the gap is largest on data reconstruction and integration. Against both MOFA and AE, we see a large improvement in topic-view quality. For FACTMx, topic-view relative log-likelihood collapses to ≈ 1 (matching the generative model) versus ≈ 1.8 for AE, and the topic component distances are about an order of magnitude smaller. This comes as a consequence of MOFA and AE using pre-summarised vectors as input rather than modelling topics directly. Against FACTM, the greatest gains are seen in data integration. FACTM’s mean view *R*^2^ stays at ≈ 0.15–0.30 across all scenarios whereas FACTMx reaches ≈ 0.85–0.92, indicating that FACTMx’s flexible variational coupling integrates simple and structured views into the latent representation far more effectively than FACTM’s tailored linear formulation. FACTMx outperforms competitors in all GMM-related metrics in most scenarios, due to its ability to infer the Gaussian components jointly with their dependencies on latent patient representations. However, we do observe one regime where FACTMx itself degrades, namely combinations of dense latent-view dependencies with substantial noise. There GMM component recovery becomes noisier, although the general model performance still matches or exceeds the competitors’. Overall, FACTMx is the only method that holds up across all data regimes and the areas where its advantage is largest, namely structured-view quality and data integration, are precisely the ones crucial in real cohort studies.

### FACTMx latent captures interpretable cross-modal survival signature in NSCLC patients

We used FACTMx to analyse a non–small cell lung cancer (NSCLC) dataset generated by the IMMUcan consortium [6], which comprises 192 patients profiled across four data modalities, including one simple: bulk RNA sequencing, and three structured: multiplex immunofluorescence (IF) imaging on three tissue sections using three antibody panels, imaging mass cytometry (IMC) on two tissue sections using two antibody panels, and hematoxylin and eosin (H&E)–stained histology on two tissue sections. We aimed to integrate these imaging and molecular contexts into a shared latent representation, then assess whether the model recovers clinically meaningful and survival-associated signals. For comparison, we also trained FACTM, an ablation AE, and MOFA models on the same data after the necessary preprocessing steps for structured modalities (see Methods).

The application workflow (Fig. 2**a**, Methods) was as follows. For structured view inputs, patient H&E images were divided into patches, which were subsequently embedded using the UNI2 model [19]; IMC tumour regions of interest (ROIs) for IMC panels 1 and 2, and tertiary lymphoid structure (TLS) regions for IMC panel 1 were segmented into cell niches, represented as celltype counts in the niche; images from all IF panels were divided into ROI (tumour) and next-to-tumour (stroma) regions, and segmented into cell niches. For simple view inputs, we used top 1000 variably expressed genes with matching HUGO [20] symbols. After training the model, we extracted the patient-level representations, patient component proportions, and the latent factor weights for analysis (see Methods). For interpretability, inferred niche types were labelled using celltypes with highest frequencies in their profiles (Supp. Figs. 1–5) and inferred H&E components were labelled in consultation with a pathologist (Supp. Figs. 6–17). Since not all 192 patients had data available across all views (Fig. 2**b**) and FACTMx does not handle missing view data, the training cohort was restricted to 102 patients. In both the analysed and full cohorts, there was a slight prevalence of male patients and stage 1 disease, a strong prevalence of the adenocarcinoma subtype, and a balanced distribution of smoking history (Fig. 2**c**, Supp. Fig. 18).

**Figure 2:**
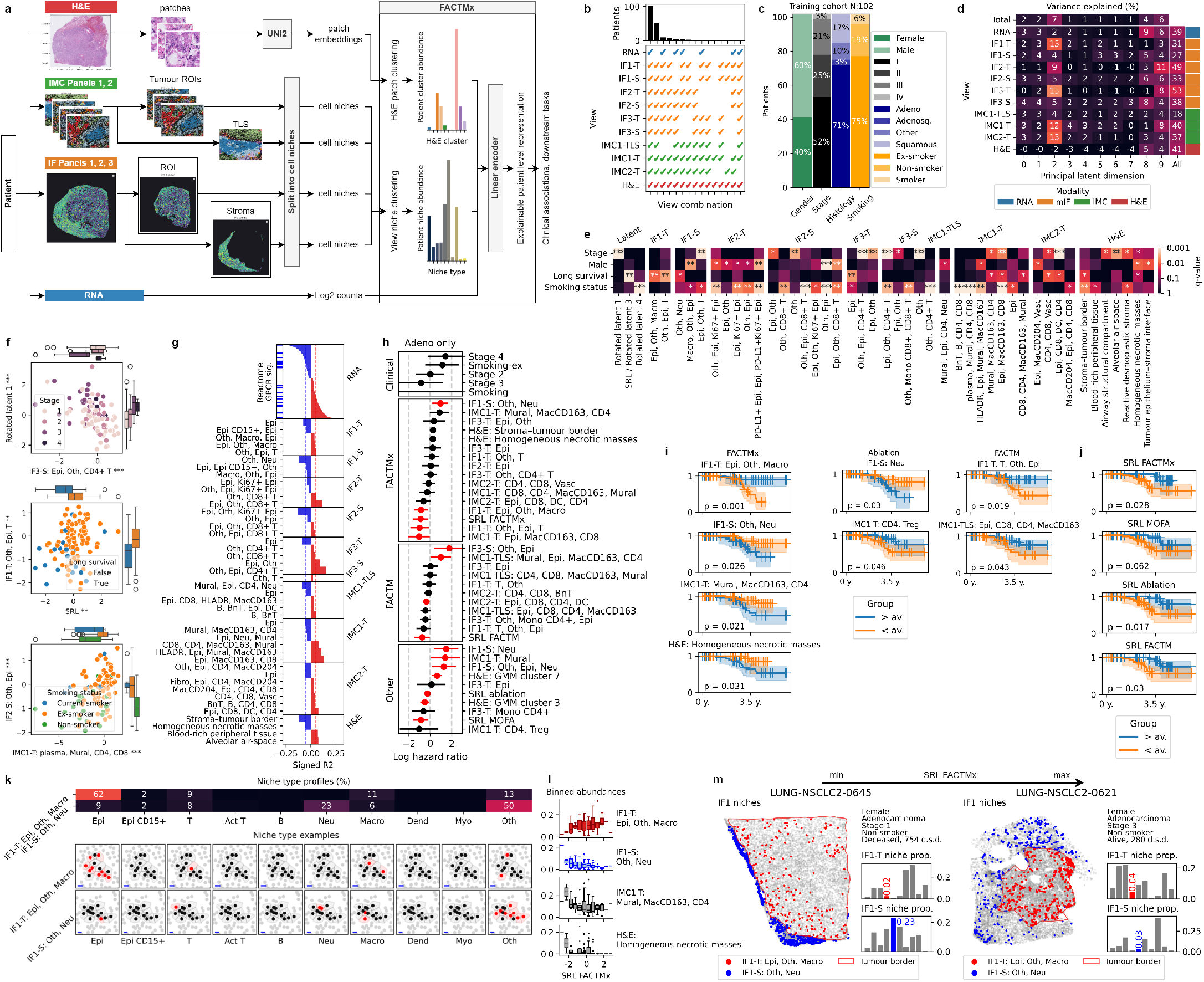
FACTMx results analysis for NSCLC patients from the IMMUcan NSCLC2 cohort: **a** FACTMx pipeline for cohort data; **b** data availability for cohort patients; **c** training cohort summary by clinical features; **d** % variance explained per view/factor pair, total data/factor, and view/all factors; **e** ANOVA F-test − log_10_ *q*-values for clinically-rotated features and structured view predictors; **f** top 2 predictors from the ANOVA F-test vs. the associated clinical variables of patients; **g** signed *R*^2^ of the long-survival-rotated factor for features with *R*^2^>0.02; **h** Cox model hazard ratios for predictors significantly associated with survival and clinical covariates for adenocarcinoma patients, 95% confidence intervals shown, statistically significant results marked in red; **i** Kaplan-Meier (K-M) curves for adenocarcinoma patients stratified by predictors significantly associated with survival; **j** K-M curves for adenocarcinoma patients stratified by SRL of different methods; **k** niche profiles and examples for top and bottom survival associated niche types; **l** average proportions of FACTMx components from **i** across FACTMx SRL; **m** patients with min and max values of FACTMx SRL with their IF1 slides. *Adeno* – adenocarcinoma, *Adenosq*. – adenosquamous, *Epi* – Epithelial, *CD4* – CD4+ T cells, *CD8* – CD8+ T cells, *Treg* – regulatory T cells, *DC/Dend* – dendritic cells, *Oth* – other cells, *Mono* – monocytes, *Neu* – neutrophils, *Mac/Macro* – macrophages, *BnT* – B and T cells, *Myo* – myeloid, *Vasc* – vasculature, *SRL* – survival rotated latent, *sig*. – signalling, *d*.*s*.*d*. – days since diagnosis, ∗ *<* 0.05, ∗∗ *<* 0.01, ∗ ∗ ∗ *<* 0.001.

As the linear latent space is invariant to rotation, it can be re-expressed for specific downstream analyses. We first used principal component analysis (PCA) on the latent representations to assess how the multimodal variation was distributed in the latent space (Fig. 2**d**). The proportions of variance explained showed one dimension shared across all views (dimension 9), a dimension shared across topic views but especially tumour (dimension 2), a dimension shared by RNA, IF stroma views and to a lesser degree IMC and H&E views (dimension 8), and seven mixed dimensions (dimensions 0-1, and 3-7). This pattern indicates that FACTMx captured a broad multimodal signal containing both shared and modality-specific structure. We then rotated the already learnt latent space towards selected clinical variables – subtype, stage, sex, long survival, smoking history, and KRAS and TP53 mutation status —- to obtain clinically-oriented factors of that broader signal for downstream interpretation. The rotated dimensions should therefore be understood as endpoint-focused factors of the discovered latent structure, rather than as independent model outputs.

Next, we used the ANOVA F-test to see which rotated latent dimensions (RLs) and patient component proportions differ significantly between patients stratified by stage, sex, long survival, or smoking history (Fig. 2**e**, *q*-values obtained with Benjamini-Hochberg (B-H) correction). The RLs showed a significant association with their rotation target covariates (except sex), and every structured view had at least one component associated to one of the covariates. We plotted the top 2 predictors of stage, survival, and smoking history against their covariates (Fig. 2**f**), visualising the combined strength of the multimodal signal.

To understand which features contribute to the survival-rotated latent dimension (SRL), we used a signed *R*^2^ metric (see Methods), which describes how well each feature of the data is reconstructed using SRL and whether it is positively or negatively loaded (Fig. 2**g**). Notable features contributing to SRL included the IF panel 1 tumour (IF1-T) niche type comprising epithelial, other, macrophage, and T cells (referred to as IF1-T: epi, oth, macro in Fig. 2**g**; here shortly EOM); and the IF panel 1 stroma (IF1-S) niche type comprising other, neutrophil, and a smaller admix of epithelial cells (IF1-S: oth, neu; ON). In the H&E structured view, we see that the stroma-tumour border and homogeneous necrotic masses components contribute to SRL negatively, and alveolar airspace and blood-rich peripheral tissue contribute positively. By performing gene set enrichment analysis (GSEA, [21]) on the signed *R*^2^ scores, we also found that the top two enriched Reactome [22] pathways contained genes connected to GPCR signalling (*ES* = 0.62, *q* = 0.038) and vesicle mediated transport (*ES* = −0.52, *q* = 0.057), the latter of which is linked to tumour growth, metastasis, and resistance in NSCLC [23–25]. Together, these results link SRL with a multimodal signature of immune response, invasive and necrotic tumour vs. healthy tissue content, and tumour growth.

We performed two types of survival analysis, limiting ourselves to adenocarcinoma patients only, as they comprised the majority of the analysed cohort. Firstly, we used the Cox proportional hazards model, which in the baseline setting included stage and smoking history information, with non-smoking stage 1 patients as reference (clinical in Fig 2**h**). Then, for each FACTMx feature that was significant in the ANOVA F-test of long-survival patients (long-survival features), we fitted a Cox model with that feature, stage, and smoking information, again keeping non-smoking stage 1 patients as reference (FACTMx in Fig 2**h**). We avoided one combined model due to the high correlation of features from different views and the SRL. The same analysis was then performed for long-survival features from competitor methods (FACTM; precomputed summaries of structured views or factors derived by ablation AE and MOFA, jointly referred to as ‘Other’ in Fig. 2**h**). FACTMx SRL showed the largest decrease in patient hazard function among latent predictors, again showcasing its stronger multimodal signal. The spatial niches IF1-T EOM and IF1-S ON also obtained significant hazard ratios. Secondly, for each long-survival feature we split the patients into those with that feature’s value larger than its cohort mean (over) and smaller than that mean (under). We then used log-rank tests to investigate whether the at-risk fractions differed significantly between the two groups for each feature, and plotted the Kaplan-Meier (K-M) curves for all structured features with significant *p*-values (Fig. 2**i**) and SRLs from all methods (Fig. 2**j**). FACTMx had more statistically significant structured predictors that better stratified patients into risk groups, with above average IF1-T EOM proportions being especially favourable for long survival. SRLs of all methods, except MOFA’s, achieved statistically significant stratification.

The interpretable structure of the FACTMx analysis provides a basis for generating hypotheses about the biological processes underlying survival-associated features. We considered the two niche types from IF panel 1 that influence the FACTMx SRL and were significant in Cox and K-M analyses: the EOM and ON niche types (Fig. 2**k**, Supp. Figs. 19–27). Their celltype frequency profiles are consistent with the hypothesis that the IF1-T EOM niche type captures microenvironments involving macrophage-mediated T-cell recruitment during an anti-tumour response [26, 27]. Similarly, the IF1-S ON niche type is consistent with dense aggregates of tumour-associated neutrophils [28, 29]. These model-derived interpretations could be tested in additional sequencing experiments, for example using markers such as CXCL9 or CD177. The link between SRL and survival-stratifying features can also be seen in how their abundances change as SRL increases (Fig. 2**l**), where IF1-T EOM and IF1-S ON show close to monotonic relationships. Finally, comparing spatial distribution of patients with the highest and lowest SRL values (Fig. 2**m**), we found that the high-SRL patient had more abundant IF1-T EOM and sparse, low levels of IF1-S ON, whereas the low-SRL patient showed the opposite pattern.

In conclusion, we showcased how FACTMx can be deployed on heterogeneous cohort-scale NSCLC data to facilitate recovery of clinically relevant latent factors, linking them to interpretable spatial and molecular features, and highlighting survival-associated structure with a signal stronger than that of the comparative analyses considered here, which was not summarised by the baseline clinical covariates.

### FACTMx analysis links spatially-coherent tumour microenvironments to survival in large-scale NSCLC cohort

We then joined the TCGA LUAD and LUSC cohorts [7, 8] to test FACTMx in a larger NSCLC cohort setting with a different combination of modalities. To define the views for simple modalities, we selected the top 1000 variably expressed genes, the top 1000 variably methylated positions, and the top 1000 frequently altered genomic positions for copy number alterations (CNAs; see Methods). Additionally, as a structured view, we segmented patient H&E slides into patches and embedded them with the UNI2 [19] model (Fig. 3**a**). After selecting patients with data across all views, the final cohort size was *N* = 881 (Fig. 3**b**). Male patients were as prevalent as in the IMMUcan cohort. However, stage 3 patients were less common, as were non-smokers. Since the LUAD and LUSC cohorts are similar in size, the joined cohort was balanced across the subtypes.

**Figure 3:**
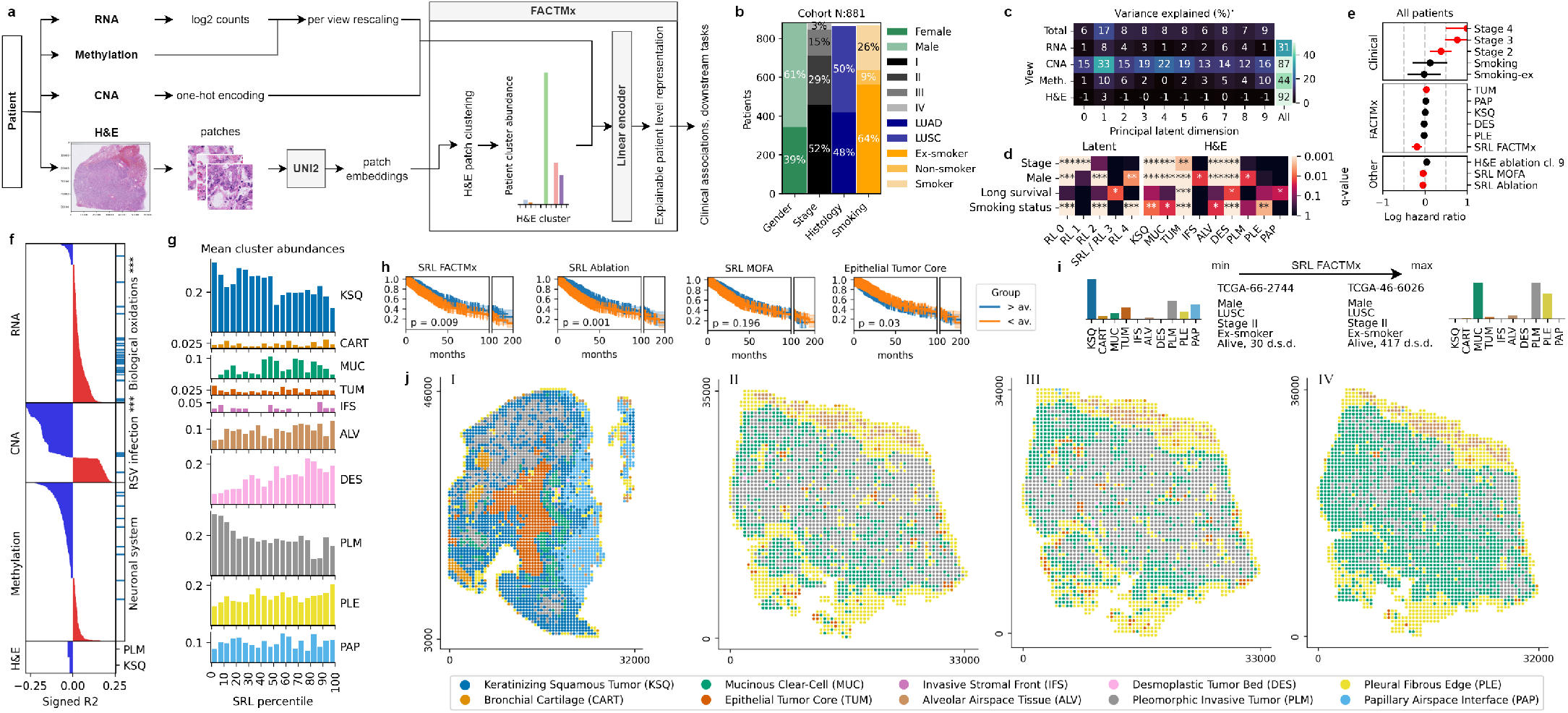
FACTMx results analysis for NSCLC patients from LUAD and LUSC TCGA cohorts: **a** FACTMx pipeline for cohort data; **b** training cohort summary by clinical features; **c** % variance explained per view/factor pair, total data/factor, and view/all factors (for categorical CNA this is instead relative likelihood); **d** ANOVA F-test − log_10_ *q*-values for clinically-rotated features and structured view predictors; **e** Cox model hazard ratios for clinical covariates and top predictors from ANOVA F-test of FACTMx and competitor methods marked as Other (MOFA, ablation AE, and precomputed patient summaries from structured views), 95% confidence intervals shown, statistically significant results marked in red; **f** signed *R*^2^ of the long-survival-rotated factor for features with *R*^2^>0.02; **g** binned cluster abundances for patient percentiles across FACTMx SRL; **h** Kaplan-Meier curves for patients stratified by predictors significantly associated with long survival; **i** comparison of patients with min and max values of FACTMx SRL; **j** H&E slides of patients with min (I) and max (II-IV) values of SRL, coloured by patch assignment in the model. ∗ *<* 0.05, ∗∗ *<* 0.01, ∗ ∗ ∗ *<* 0.001, *(S)RL* – (survival) rotated latent, *d*.*s*.*d*. – days since diagnosis.

For latent-space analysis, we again used PCA and post hoc rotation. The per-view *R*^2^ scores showed that the latent representation was dominated by CNA and H&E views (Fig. 3**c**). Individual principal latent dimensions did not achieve high *R*^2^ values for the H&E component proportions, likely because these are decoded as softmax-transformed proportions, which limits their exact decoupling in the latent. Compared with the IMMUcan analysis, fewer dimensions were view-specific and the per-factor *R*^2^ values were more evenly distributed across views. This indicates that FACTMx recovered a broad cross-modal signal spanning genomic and histological structure. We then rotated the latent towards subtype, stage, sex, long survival, and smoking history to obtain clinically-oriented factors for downstream analyses (see Methods). As above, these rotated latent (RL) dimensions should be interpreted as analysis-specific views of the broader learnt representation.

We used the ANOVA F-test to see which RLs and patient H&E component proportions differed significantly between patients stratified by stage, sex, long survival, or smoking history (Fig. 3**d**, *q*-values obtained with B-H correction). The RLs showed a significant association with their target clinical covariates. Additionally, the subtype rotated RL0 showed significant differences in stage-, sex-, and smoking-stratified patients. The most notable H&E component in this analysis was epithelial tumour core (TUM), which was most significantly associated with long survival, and showed associations with all other clinical features. Stage and male sex showed associations with multiple components, including keratinizing squamous tumour (KSQ), mucinous clear cell (MUC), TUM, alveolar airspace tissue (ALV), and desmoplastic tumour bed (DES).

Next, we used the Cox proportional hazards model, which in the baseline setting included stage and smoking history information, with non-smoking stage 1 patients as a reference (clinical in Fig 3**e**). Then, for each long-survival feature (defined as in the IMMUcan analysis) from the analysed integration methods, we fitted a Cox model with that feature, stage, and smoking information, again keeping non-smoking stage 1 patients as a reference (FACTMx; precomputed summaries of structured views or factors derived by ablation AE and MOFA, jointly referred to as ‘Other’ in Fig. 3**e**). TUM was the only H&E component with a statistically significant hazard ratio. In addition, FACTMx SRL invoked the largest decrease in the patient hazard function, showcasing the strengthened multimodal signal.

To analyse which features were tied to the FACTMx SRL, we again used the signed *R*^2^ metric (Fig. 3**f**). Two H&E components explained by SRL were the pleomorphic invasive tumour (PLM) and keratinizing squamous tumour (KSQ), both with a negative effect. We also performed GSEA on the signed *R*^2^ scores from simple views using Reactome pathway gene sets, with biological oxidations pathway being the most enriched in the RNA view and the pathway tied to RSV infection being most enriched in the CNA view. The methylation view showed no pathway with statistically significant enrichment, despite having a range of features contributing to the SRL. Overall, the number of features that contribute to the SRL in all the views indicates the multimodal nature of the survival signal.

To further investigate the link between SRL and the H&E components, we analysed how their abundance changes in groups of patients with increasing SRL, where we again see SRL’s negative correlation with PLM and KSQ (Fig. 3**g**). The TUM abundance did not appear to be associated with SRL, showcasing that some orthogonal survival signal is present in the model outside of the SRL.

We then used the same K-M analysis setup as in the IMMUcan analysis for SRL and long-survival features of all methods. TUM was again the only H&E component with a statistically significant result (Fig. 3**h**). The FACTMx and ablation SRLs were comparable, with FACTMx having a better separation of patients with very long survival (100+ months).

Lastly, we compared the two patients with the highest and lowest FACTMx SRL values (Fig. 3**i-j**). On their slides, the spatial distributions of H&E components were highly coherent, even across the three consecutive tissue slices of the top patient (II-IV in Fig. 3**j**), highlighting the biological consistency of the FACTMx clustering.

To summarise, we demonstrate that FACTMx can be used to analyse patient cohort data, integrating heterogeneous NSCLC subtypes across proteomic, genomic, and histopathological views. This integration allows the recovery of a strong survival-associated signal as evidenced by the ANOVA, Cox, and K-M analysis, which can be analysed and interpreted directly. Additionally, the inferred clustering of the H&E modality shows high spatial coherence and supplements TUM as an orthogonal survival associated feature.

### FACTMx factors linearly model outcome-linked immune signature in coronary syndrome patients

Next, we applied FACTMx to a longitudinal multimodal cardiovascular cohort comprising coronary syndrome patients. After modality-specific quality control and rescaling, the model integrated peripheral blood mononuclear cell (PBMC) scRNA-seq, plasma cytokine (CK) measurements, plasma proteomics, polymorphonuclear neutrophil (PMN) Prime-seq profiles, and clinical blood variables into a shared latent space, from which patient representations were extracted for downstream clinical association analyses (Fig. 4**a**, Methods). The cohort contained both acute coronary syndrome (ACS) and chronic coronary syndrome (CCS) patients (Fig. 4**b**), with repeated sampling across multiple ACS timepoints (Fig. 4**c**), providing a setting in which it was possible to test whether FACTMx could jointly capture disease state, temporal recovery and clinical outcome from heterogeneous measurements.

**Figure 4:**
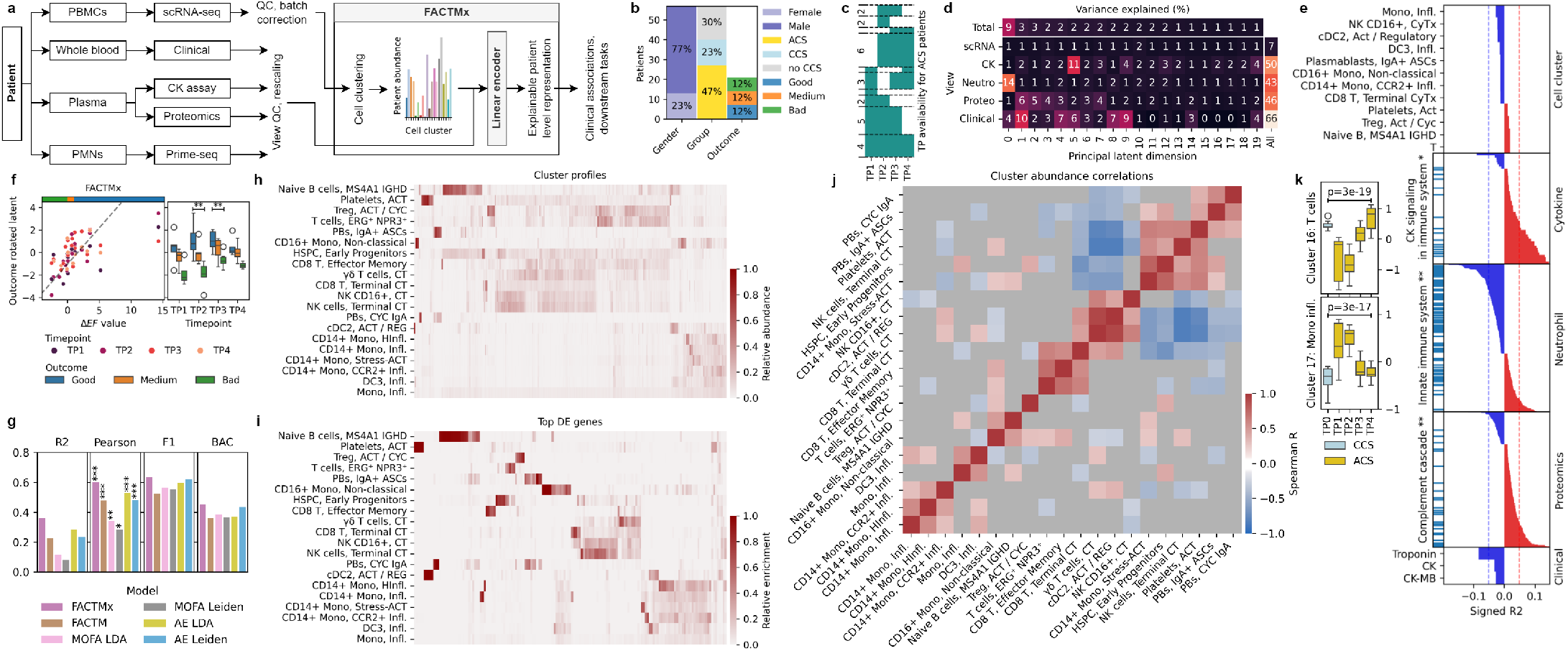
FACTMx results analysis for ACS/CCS patients: **a** FACTMx pipeline for ACS/CCS data; **b** clinical summary of the patients; **c** summary of timepoint sample availability for ACS patients; **d** % variance explained per view/factor pair, total data/factor, and view/all factors; **e** signed *R*^2^ of the outcome-rotated factor for features with *R*^2^>0.01, with most statistically enriched pathways shown for cytokine, neutrophil, and proteomics views, with marked gene/protein positions and *q*-values; **f** outcome-rotated factor vs. Δ*EF* and time from acute event, with trendline and ANOVA *p*-values; **g** evaluation of outcome-rotated factors outcome prediction by *R*^2^ of linear regression, Pearson’s R, and weighted F1 score and adjusted balanced accuracy of a logistic regression; **h** relative abundance of gene probabilities in profiles of FACTMx clusters; **i** relative enrichment of log-foldchanges in all gene expression of FACTMx clusters; **j** correlations of FACTMx cluster abundances in patients; **k** log transformed relative abundances of cell clusters best explained by FACTMx outcome-rotated factor, by timepoint. *CK* – creatine kinase, *neutro* – neutrophil, *proteo* – proteomics, *TP* – timepoint, *mono* – monocytes, *CyTx/CT* – cytotoxic, *Act* – activated, *Infl*. – inflammatory, *HInfl*. – hyperinflammatory, *Cyc* – cycling, *PBs* – plasmablasts, *AE* – ablation autoencoder, ∗ *<* 0.05, ∗∗ *<* 0.01, ∗ ∗ ∗ *<* 0.001.

As before, we used PCA and post hoc rotation for latent analysis. Latent PCA showed distribution of variance across all views, indicating that the model was not dominated by a single assay but instead captured both shared and view-specific structure (Fig. 4**d**). Clinical variables, proteomics, and neutrophil-derived measurements contributed strongly to the overall variance explained, while scRNA-seq and cytokine features added complementary signal across multiple latent dimensions. This indicates that FACTMx recovered a broad multimodal representation of disease state and recovery. For downstream interpretation, we rotated the latent towards outcome (difference in ejection fraction, Δ*EF*), sex, classification, and age to obtain clinically-oriented factors, which should be interpreted as endpoint-specific views of the broader learnt structure (see Methods).

Feature-level analysis of the outcome-rotated latent factor (ORL) using the signed *R*^2^ metric highlighted its multimodal nature (Fig. 4**e**). Among cellular component features, the negative associations involved levels of inflammatory monocyte, cytotoxic NK, and CD8 T-cells. Neutrophil features from the innate immune system pathway showed negative enrichment. Plasma CK and proteomics features positively associated with ORL were linked with immune system signalling and complement cascade biology respectively, while classical markers of myocardial injury, including troponin, CK and CK-MB, were negatively associated. These associations link FACTMx ORL to a combination of systemic inflammation, innate and adaptive immune activation, and biochemical evidence of cardiac damage. The ORL captured a linear dependency with ΔEF and separated patients according to outcome class, with clearer stratification at follow-up timepoints after the acute event (Fig. 4**f**). These results link ORL to possible differences in immune state during recovery after acute injury.

We compared FACTMx ORL with outcome-rotated latent variables obtained using alternative integration strategies: FACTM, MOFA, or ablation AE. The latter two had two variants, which used precomputed summaries of the cellular component structured view obtained with either LDA or Leiden clustering. Across regression and classification benchmarks, FACTMx consistently performed best, yielding the highest *R*^2^, Pearson correlation, weighted F1 score, and adjusted balanced accuracy (Fig. 4**g**). This improvement suggests that joint modelling of patient representations using FACTMx improves the recovery of clinically relevant structure beyond standard factorisation or autoencoder-based embeddings.

To interpret the cellular component structured view of the model, we examined the inferred corresponding component profiles. FACTMx identified coherent cell-state profiles encompassing naïve and activated B-cell populations, cytotoxic and terminal T/NK states, dendritic-cell subsets, plasmablast/ASC populations, and multiple monocyte inflammatory states (Fig. 4**h**). Differential gene expression between these clusters supported their biological distinctiveness and reinforced the interpretability of FACTMx latent space in terms of discrete but related immune programmes (Fig. 4**i**). Correlation analysis of patient cluster abundances revealed positively correlated modules of related cell states and anticorrelations between inflammatory myeloid and lymphoid compartments, indicating coordinated remodelling (Fig. 4**j**).

Finally, we examined cell populations most strongly linked with ORL. The abundances of T-cell cluster 16 and inflammatory monocyte cluster 17 varied significantly across ACS timepoints, while there was comparably less variation within CCS samples (Fig. 4**k**). The inflammatory monocyte population was highest early after the acute event and declined over follow-up, while the T-cell-associated cluster showed the opposite tendency. These temporal changes further the interpretation that FACTMx ORL captures a transition from an early innate inflammatory phase towards a later, more resolved immune state associated with a more favourable cardiac recovery.

Overall, these analyses show that FACTMx can integrate longitudinal single-cell, plasma, neutrophil and clinical measurements into a unified patient representation that is both interpretable and clinically informative. In the ACS/CCS cohort, model analysis recovered an outcome-associated latent axis linked to ΔEF, was associated with differences between favourable and unfavourable recovery trajectories, and connected these trajectories to coordinated changes in inflammatory monocytes, lymphocyte states, soluble immune mediators and cardiac injury markers.

## Discussion

In this work, we present FACTMx, a flexible variational framework for integrating simple and structured views into explainable patient-level representations. In contrast to approaches that rely on fixed summaries of structured views obtained in a separate preprocessing step, FACTMx models lower-level structure jointly with the latent patient representation. This improves both the quality of integration and the correspondence between patient-level factors and structured observations. Across simulated and experimental datasets, FACTMx outperformed comparative methods according to validation metrics and the strength of recovered clinical associations, respectively. The model formulation supports the interpretation of downstream analyses in clinically relevant terms. For the IMMUcan NSCLC cohort, FACTMx analysis highlighted a survival-associated signal that could be related to spatially resolved immune microenvironments, interpretable H&E components, and known pathways of tumour growth and metastasis. In the TCGA lung cancer cohorts, the same analysis strategy linked a survival-oriented view of the latent space to coherent histological patterns in H&E slides. In the coronary syndrome cohort, it yielded an outcome-oriented view of the latent space connected to temporal changes in cellular and molecular measurements. Together, these results suggest that FACTMx excels in learning patient-level multimodal representations and organising them into clinically interpretable analytical factors for downstream study.

Several limitations should also be noted. First, the modest size of currently available multimodal cohorts does not support the full nonlinear FACTMx configuration, prohibiting us from exploring settings where its highly expressive parameterisations may be more advantageous. As larger and more complete multimodal cohorts become available, the benefit of flexible nonlinear dependencies should become easier to realise in practice. On the other hand, with the current linear parametrisation, the model enjoys higher interpretability. Second, the quality of the learnt representation still depends on hyperparameter choice, including the assumed latent dimensionality, per-modality loss scales, and head-specific parameters. Missingness across modalities, which is common in real cohorts, may reduce the amount of information available for joint estimation. Lastly, the present implementation focuses on mixture-type structured views, but the general FACTMx formulation is not limited to this case. Extensions to other classes of structured latent-variable models should therefore be possible without changing the overall modelling principle and with minimal implementation cost.

Despite these limitations, FACTMx already proves a valuable tool addressing the emerging need of integrating structured and simple views in multimodal patient profiling cohorts, providing interpretable patient-level representations and allowing for deep analyses of inter-modal statistical signals.

## Methods

**IMMUcan NSCLC data**

We analysed a non–small cell lung cancer (NSCLC) dataset generated by the IMMUcan consortium [6]. The dataset comprises 192 patients profiled across four data modalities: bulk RNA sequencing, multiplex immunofluorescence (IF) imaging on three tissue sections using three antibody panels, imaging mass cytometry (IMC) on two tissue sections using two antibody panels, and hematoxylin and eosin (H&E)–stained histology on two tissue sections. Sample availability across modalities is summarised in Supp. File 1.

Bulk RNA-seq libraries were prepared from FFPE or frozen tissue using either the KAPA RNA HyperPrep Kit with RiboErase (H/M/R) Globin (Roche) or the SMART-Seq Stranded Kit (Takara Bio), depending on RNA quality and quantity. Sequencing was performed on an Illumina NovaSeq 6000 platform. Gene expression data were filtered for minimal expression, adjusted for sequencing kit, library-size normalised, and log1p-transformed. We kept top 1000 most variably expressed genes annotated with HUGO symbols. To harmonise variance across modalities while preserving relative gene-level variability, expression values were scaled so that the mean gene-wise variance across the cohort equalled 1.

Multiplex IF data were generated from three 4-µm FFPE sections using three antibody panels as described by Schultz *et al*. [6]. Segmented IF images were partitioned into tumour and next-to-tumour regions by a pathologist. IF cellular niches were obtained by using a BallTree scikit-learn [30] implementation with leaf size 15, followed by clustering of leaf nodes into connected components with connectivity radius 15*µ*m; a single representative cluster from each leaf node was retained as a niche if its cell count fell between 6 and 20 cells. Niche compositions were defined as their celltype counts.

IMC data were generated under IMMUcan specifications [31] from two tissue sections using two antibody panels. Cell-level data from tumour and tertiary lymphoid structure (TLS) regions were grouped into spatial niches using hierarchical clustering of cell coordinates, using Ward linkage and a distance threshold of 50*µm*; then, niche compositions were defined as their celltype counts.

H&E-stained slides were processed using the Trident pipeline [32] with 20× magnification and 256×256 pixel patches. Patch embeddings were extracted using the UNI2 model [19] (1,526-dimensional embeddings), reduced by PCA to 50 components and rescaled by 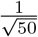.

### TCGA NSCLC data

We analysed lung adenocarcinoma (LUAD; TCGA-LUAD) and lung squamous cell carcinoma (LUSC; TCGA-LUSC) cohorts from The Cancer Genome Atlas (TCGA) accessed via the Genomic Data Commons (GDC). The data are part of the TCGA PanCancer Atlas (PanCanAtlas) initiative, which provides harmonized multi-omic profiles for 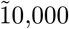 tumours across 33 cancer types, including updated and expanded LUAD and LUSC cohorts relative to earlier studies [7, 8, 33]. The PanCancer Atlas represents a unified set of molecular and clinical data processed with consistent pipelines, including RNA sequencing, exome sequencing, and DNA methylation assays, across all patients.

Bulk RNA sequencing was performed by TCGA using poly(A)-selected libraries prepared with the Illumina TruSeq RNA Sample Preparation Kit and sequenced on Illumina platforms as part of the PanCancer Atlas data production pipeline [34]. In our analysis, we included the 1,000 most variably expressed genes across patients.

Whole-exome sequencing data were generated using Agilent SureSelect Human All Exon capture kits and sequenced on Illumina platforms following TCGA protocols [35]. Copy number alteration (CNA) profiles derived from these data were encoded as one-hot vectors across five states (−2, −1, 0, +1, +2), with the 1,000 most frequently altered genomic positions across patients retained.

Genome-wide DNA methylation was assayed on the Illumina HumanMethylation450 BeadChip array according to TCGA procedures [36]. We selected the 1,000 most variably methylated positions across patients.

H&E whole-slide images were processed using the Trident pipeline with 20× magnification and 256×256 pixel patches. Patch-level features were extracted with the UNI2 model (1,526-dimensional embeddings), reduced by PCA to 100 components, and rescaled by 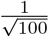.

Additionally, all simple views were rescaled such that the mean variance of their features across the cohort equalled 1.

### ACS/CCS data

We analysed a cohort of patients with acute coronary syndrome (ACS) and chronic coronary syndrome (CCS), originally described in [5]. The cohort comprises 62 patients, with ACS patients sampled at up to four time points following the acute event. Five data modalities were available: scRNA-seq of peripheral blood mononuclear cells (PBMCs), plasma cytokine profiling, plasma proteomics, bulk neutrophil transcriptomics, and routine clinical blood tests. Sample availability across modalities and time points is summarised in Supp. File 2.

single-cell RNA-seq libraries were generated from frozen PBMCs using the Chromium Next GEM Single Cell 3’ Reagent Kit v3.1 (10x Genomics), with barcode-based multiplexing using TotalSeq-B anti-human hashtag antibodies. Preprocessed scRNA-seq data from the original study (following demultiplexing, doublet removal, and quality filtering) were filtered to the top 2,000 highly variable genes per library and merged using the 417 genes identified as highly variable across all libraries. Batch correction and joint embeddings were obtained using Scanorama [37] with default parameters.

Plasma cytokine and chemokine concentrations were measured using a Human Cytokine/Chemokine 71-Plex Discovery Assay (Eve Technologies). Values outside the detection range were set to zero, analytes detected in at least 20% of samples were retained, and data were log1p-transformed. After filtering, cytokine data was available for 126 samples across 71 features.

Bulk neutrophil transcriptomes were generated using prime-seq. Expression data were filtered to retain genes detected in more than 80% of patients, excluding mitochondrial and ribosomal genes; patients with more than 25% missing values were excluded. Gene expression values were mean-scaled and log1p-transformed. In a final filtering pass, genes with variance lower than the 25th quantile of all gene-wise variances were excluded. After filtering, neutrophil data was available for 118 samples across 892 features.

Plasma proteomics data were acquired using label-free data-independent acquisition mass spectrometry on an Orbitrap Exploris 480 instrument. Data preprocessing followed the pipeline described in the original publication, followed by a final centering. Plasma proteomics data was available for 119 samples across 490 features.

Clinical blood test data were obtained as part of routine clinical care during hospitalisation and included creatine kinase (CK), CK-MB, troponin T, and C-reactive protein (CRP). Missing or out-of-range values were set to zero, followed by log1p transformation and normalisation.

Additionally, all simple views were scaled such that the mean variance across features in the view equalled 1.

Clinical outcome was defined based on serial measurements of left ventricular ejection fraction (EF), assessed by echocardiography using Simpson’s method. The change in EF (ΔEF) was computed as the difference between the value recorded on admission and either the first in-hospital measurement or the value obtained prior to discharge. Patients were classified as having a good outcome if ΔEF was greater than 1, medium outcome if 0 ≤ ΔEF ≤ 1, and a bad outcome if ΔEF was negative.

### Simulated data

To test implementation correctness and evaluate model performance, we simulated datasets with known ground truth using the generative model underlying FACTMx. Importantly, this model satisfies the assumptions of FACTM and MOFA, which we use as comparison. We benchmarked FACTMx performance against FACTM, MOFA, and a simple autoencoder ablation model.

The simulation procedure can be summarised as follows: sample patient latent representations *z*_*i*_; for each simple modality *s*, multiply *z*_*i*_ by a decoding matrix *W*_*s*_ to obtain the parameters of that modality’s data distribution, and generate simple view data using these decoded parameters; for each structured modality *k* multiply *z*_*i*_ by *W*_*k*_ to get the unnormalised log proportions of that modality’s components for patient *i*; assign each subobservation for that patient to a component using these proportions, and generate subobservation data using the component parameters and assignments.The detailed algorithm description is provided in the Supplementary Methods.

### Preprocessing input to comparative methods

Since only FACTMx can handle collections of subobservations per patient as input data for all types of data distributions, we summarised these collections into per-patient vectors for use as input in comparative models. To do this, we used three methods: Latent Dirichlet Allocation (LDA), Gaussian mixture models (GMM), and Leiden clustering.

We used LDA to summarise the IMMUcan IF and IMC views, the single-cell data in the ACS/CCS dataset, and the topic modalities in the simulated data. For IMMUcan data, for each IF region-panel combination, we trained LDA models with 10 components on 40,000 niche compositions randomly subsampled from the full patient cohort, where each composition is a vector of celltype counts within a single niche. The trained model was then applied to all niches of each patient to obtain per-niche component assignments; each patient was subsequently summarised as the vector of proportions of their niches assigned to each component. This produced six vector summaries per patient corresponding to the three panels and the tumour and next-to-tumour regions. Similarly, IMC summaries were obtained by fitting LDA models (10 components) for each region–panel combination, yielding three summaries per patient, defined by component proportions for IMC panel 1 tumour, IMC panel 1 TLS, and IMC panel 2 tumour. For ACS/CCS single-cell data, we fit a 20 component LDA to the batch-corrected count matrix and obtained a per-patient summary by taking the proportion of each patient’s cells assigned to each component. For the topic modalities in the simulated data, we fitted a 10 component LDA to 10,000 subsampled subobservations from all patients and obtained a per-patient summary by taking the proportion of each patient’s cells assigned to the found components.

We used GMM to summarise the H&E views in the IMMUcan and TCGA datasets and the GMM modalities in the simulated data. In both the IMMUcan and TCGA datasets, patient H&E summaries were obtained by fitting a GMM with 10 components to the PCA-reduced patch embeddings and summarising each patient as proportions of their patch assignments. For simulated GMM modalities, we subsampled 10,000 subobservations from all patients, initialised 10 component means with Birch clustering, and fit a 10 component GMM model. Finally, per-patient summaries were obtained by taking the proportion of each patient’s patches assigned to each component.

We used Leiden clustering as an alternative method for summarising single-cell data in the ACS/CCS dataset. We used the scanpy implementation with parameters (n_neighbors = 10, n_pcs = 50, resolution=1, random_state=0, flavor=‘igraph’, directed=False) applied to the Scanorama cell embeddings. Per-patient summaries were obtained by taking the proportion of each patient’s cells assigned to each cluster.

### The FACTMx model

Here we present an overview of the used FACTMx model setup, visualised in Figure 5. The detailed description and a more general formulation can be found in the Supplementary Methods.

**Figure 5:**
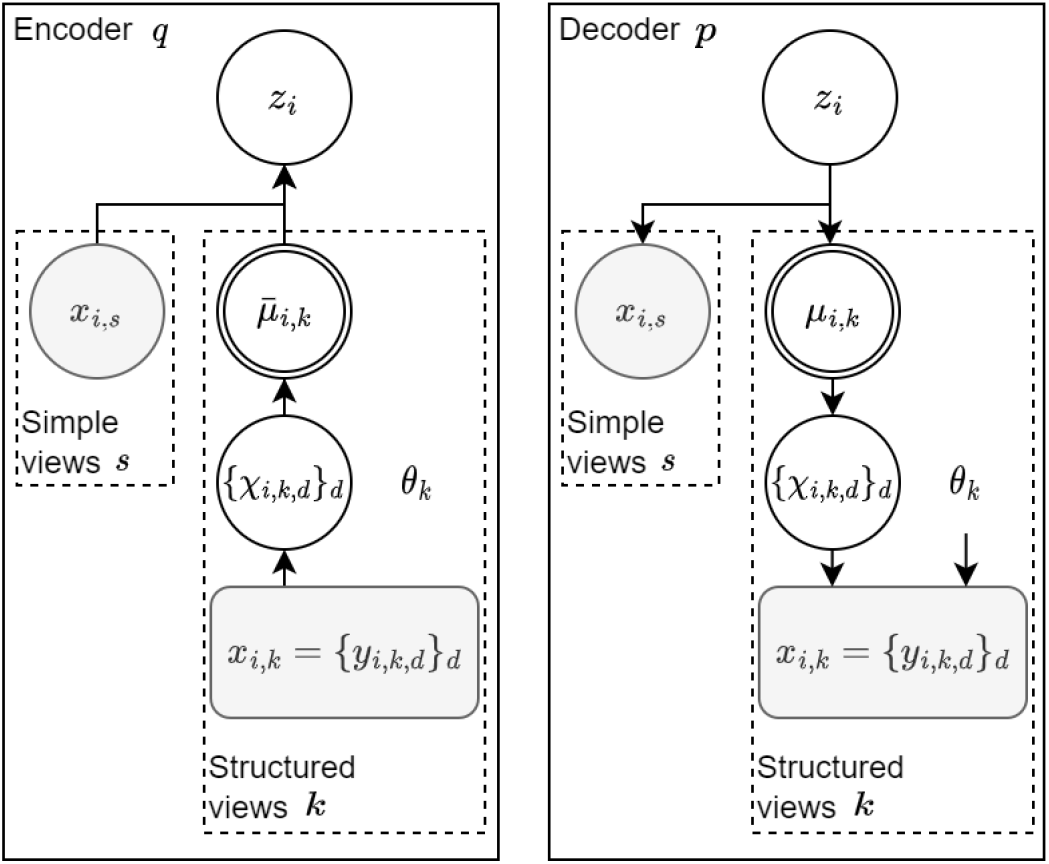
General graph formulation of the FACTMx model setup used. White nodes represent hidden variables, grey nodes represent observed variables, double-outline nodes are deterministically calculated from their parents, no-outline nodes are directly optimisable parameters.

FACTMx is a variational autoencoder combining any number of simple or structured heads that model the chosen data modalities. Simple heads represent data that can be encoded as one observed variable, while structured heads model data with distributions that factorise into bigger probabilistic graphical models (PGMs); in this publication, we focus on structured heads representing mixture models.

Formally, let *z*_*i*_ ∈ ℝ^*L*^ be an *L*-dimensional latent representation vector for sample *i*, with a prior distribution *N*(0, *I*_*L*_). Let *s, k* be indices of simple and structured views respectively. We assume each sample *i* has observations *x*_*i,s*_ and *x*_*i,k*_ that are governed by the distributions of their respective heads, with *x*_*i,k*_ being collections of subobservation data *y*_*i,k,d*_, i.e. *x*_*i,k*_ = {*y*_*i,k,d*_}_*d*_. Let *χ*_*i,k,d*_ be assignment variable for subobservation *d* of sample *i* in structured view *k*, i.e. categorical variable indicating which mixture component (topic, Gaussian, etc.) *y*_*i,k,d*_ belongs to. Let *µ*_*i,k*_ = *f*_*k*_(*z*_*i*_) be a priori proportions of components in structured view *k* of sample *i* that we assume are obtainable from *z*_*i*_ using some parametrisable function *f*_*k*_. Then *χ*_*i,k,d*_ and *µ*_*i,k*_ act as a bridge between the sample-level representation *z*_*i*_ and the raw subobservation data *y*_*i,k,d*_. Lastly, denote by *θ*_*k*_ the directly optimisable parameters of head *k*. For notation brevity, we omit indices in cases where we want to indicate a set over the omitted index; for example, *z* would indicate the set of latent representations of all samples.

Every FACTMx model can be represented as an encoder-decoder pair of PGMs. The decoder PGM approximates *p*, the generative distribution underlying the data. For each sample *i*, we assume that the full distribution *p* factorises as:

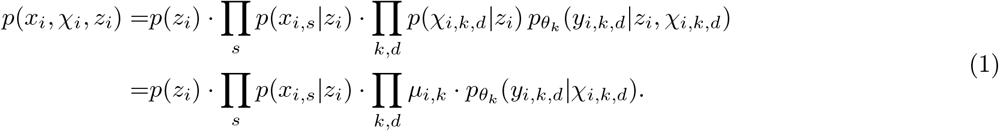

Since the posterior distribution *p*(*z*_*i*_, *χ*_*i*_|*x*_*i*_) is intractable, we use the encoder PGM to approximate it with a variational distribution *q*. It is formulated so that for each sample *i* its complete distribution factorises as:

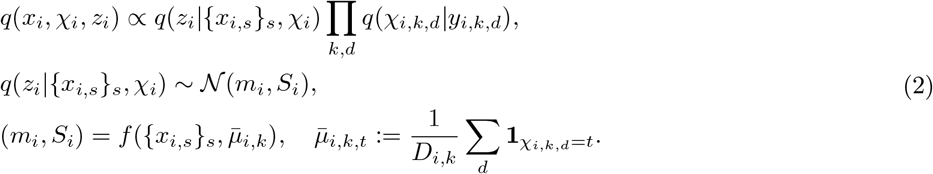

where *m*_*i*_ is the mean and *S*_*i*_ is the covariance matrix of the variational posterior, both computed by a parametrisable function *f*, taking as input the simple-view observations *x*_*i,s*_ and 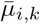, the summaries of the structured-view assignment variables *χ*_*i,k,d*_. We parametrise *q*(*χ*_*i,k,d*_|*y*_*i,k,d*_) by a deep neural classifier that takes as input subobservation data and outputs assignment logits.

To infer the parameters of *p* and *q* we use the evidence lower bound (ELBO, ℒ) objective. Due to conditional independence of views in *p* given *z*, the ELBO for sample *i* factorises as:

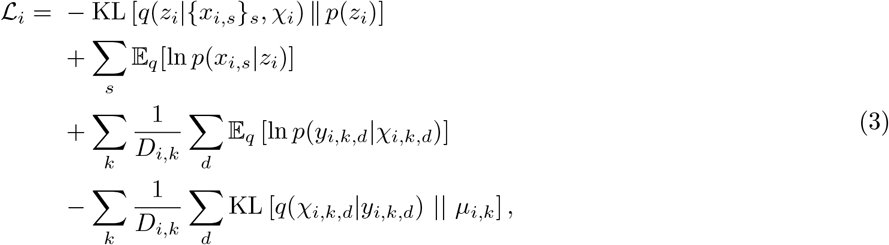

where the total ELBO is ℒ = ∑_*i*_ ℒ_*i*_.

Gradients of ℒ_*i*_ are approximated through the VAE gradient procedure (assuming reparametrisation trick):

1. For each structured view *k* and subobservation *d*, sample 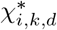 from *q*(*χ*_*i,k,d*_|*y*_*i,k,d*_).
2. Sample 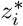 from 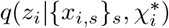.
3. For each structured view *k*, calculate 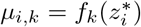
4. Calculate

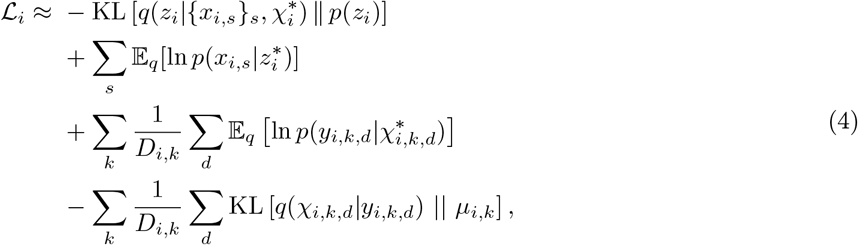
5. Backpropagate.

#### Used configurations

The following FACTMx configurations were used in this paper. In all configurations, the loss components per view were scaled by 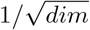, where *dim* is the data dimensionality of each view, and the encoder loss was scaled by 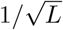.

For the IMMUcan NSCLC dataset we used a model with a linear latent encoder, linear decoders for every view, linear classifiers for topic model views, and a quadratic classifier for the Gaussian mixture view.

For the TCGA NSCLC dataset we used a model with a linear latent encoder, linear decoders for every view, and a quadratic classifier for the Gaussian mixture view.

For the ACS/CCS dataset we used a model with a linear latent encoder, linear decoders for every view, and a linear classifier for the topic single-cell view.

For the simulated datasets we used a model with a linear latent encoder, linear decoders for every view, linear classifiers for topic model views, and a quadratic classifier for the Gaussian mixture view.

#### Inference details

The FACTMx models were inferred with stochastic gradient descent using an AdamW optimiser [38] (learning rate 0.001 and weight decay 0.004).

For the IMMUcan NSCLC cohort, the model was run for 4000 epochs. In each epoch, 400 H&E patches and cellular neighbourhoods were bootstrapped per patient per view and 10 passes were made over this subsampled dataset with patient batch size 10. For the topic views, topic log-profiles were initialised from a random normal distribution with *σ* = 0.05. For the H&E view, GMM component centres and covariance matrices were initialized with random orthonormal vectors and 0.1*I* respectively.

For the ACS/CCS cohort, the model was run for 5000 epochs with patient batch size 10. In each epoch, 400 cells were bootstrapped per patient. Topic log-profiles were initialised from a random normal distribution with *σ* = 0.02.

For the TCGA NSCLC cohort, the model was run for 8000 epochs. In each epoch, 300 patients were subsampled from the whole cohort, with 200 H&E subobservations bootstrapped per patient, and 5 passes were made over this subsampled dataset with patient batch size 10. GMM component centres and covariance matrices were initialized with random orthonormal vectors and 0.1*I* respectively.

### Other methods

#### MOFA

For comparison, we ran MOFA [39] on all experimental and simulated datasets. For each dataset, we used the input matrices for the FACTMx simple views alongside the matrices with summary vectors from preprocessing of structured views. We used the implementation from mofapy2 v0.7.2 with seed = 0, dropR2 = −np.inf, keeping all other parameters default.

#### FACTM

We ran FACTM [18] on the IMMUcan, ACS/CCS, and the simulated datasets. For each dataset we used the input matrices for the FACTMx simple heads, the collections of subobservations for the FACTMx topic heads, and the matrices with summary vectors from preprocessing of the GMM views (if present). We used the original implementation from [18], allowing the model to run for 500 inference iterations, with the resulting models showing strong convergence (relative ELBO difference *<* 10^−8^).

#### Ablation autoencoder

For comparison, we implemented a simple autoencoder that concatenates all input data, passes it through a latent space with a linear encoder-decoder setup and optimises a root mean squared error (RMSE) loss during training.

The ablation autoencoder was applied to all experimental and simulated datasets, using the same input as the MOFA model. It was trained with an AdamW optimiser (learning rate 0.001, weight decay 0.004) for 500 epochs (1000 for TCGA dataset) and batch size 20.

### Analysis methods

#### Results extraction

After model inference, each sample *i* was passed through the encoder, which outputs the parameters (*m*_*i*_, *S*_*i*_) of the variational posterior *q*(*z*_*i*_ | {*x*_*i,s*_}_*s*_, *χ*_*i*_) = *N* (*m*_*i*_, *S*_*i*_); then, the mean *m*_*i*_ was taken as the patient latent representation *z*_*i*_, replacing the stochastic samples used during training. For each patient, the decoder was then applied to reconstruct the simple view data and the decoded component proportions *µ*_*i,k*_, and the assignment for all patient subobservations was sampled from *p*(*χ*_*i,k,d*_ | *z*_*i*_, *y*_*i,k,d*_).

Next, the a posteriori component proportions for structured view *k* of patient *i* were computed as the mean posterior assignments in the decoder model *p*:

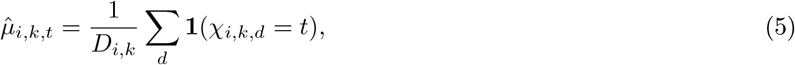

where *D*_*i,k*_ is the total number of subobservations of patient *i* in view *k* and *t* indexes the mixture components.

As input into downstream analyses, we used a log-transformation of the mean-scaled patient component proportions:

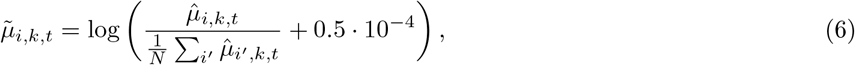

where the small constant 0.5 · 10^−4^ is added for numerical stability.

#### Per view/factor *R*^2^

To assess the influence of each latent factor *l* on each view, we first define the single-factor reconstruction of feature *j* in simple view *s* as 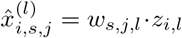, and of component proportion *t* in structured view *k* as 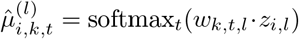, where *w*_*s,j,l*_ and *w*_*k,t,l*_ are the corresponding decoder weights. We then compute the fraction of variance explained:

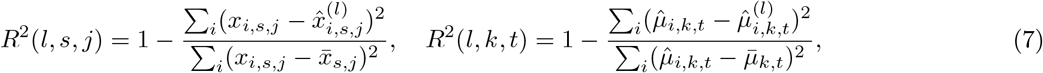

where 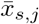 and 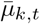 are the cohort means, and 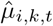 are the a posteriori proportions from Equation 5, used as reference since true proportions are unavailable. The per-factor, per-modality *R*^2^ is then:

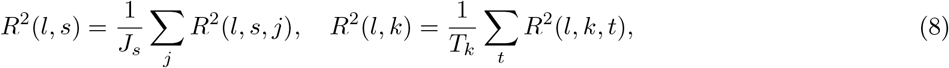

where *J*_*s*_ is the number of features in simple view *s* and *T*_*k*_ is the number of components in structured view *k*. The overall per-factor *R*^2^ is the average across all modalities and their predictors:

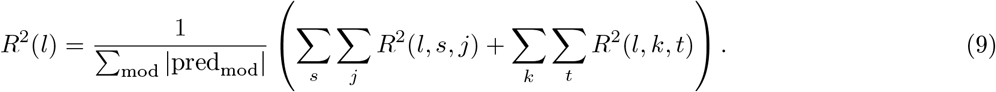

Finally, *R*^2^(mod) is computed analogously to the per-modality scores above, but using the full latent representation *z*_*i*_ rather than a single factor.

#### Rotation

For the analysis of the latent space, we used the Kabsch algorithm. In brief, up to *L* covariates are chosen, where *L* is the dimensionality of the latent space. Then, a rotation matrix *A* is calculated, such that after rotation the *l*-th dimension of the patients latent representation has the highest correlation with their *l*-th covariate. Spearman’s R was chosen as the correlation metric.

This rotation is an analysis step applied after model fitting, not a part of model training. In training, FACTMx learns a broad patient-level representation from the multimodal data without using rotation target labels in the objective. The subsequent rotation does not add new information to that representation; instead, it decomposes it into target-specific directions that are easier to interpret for a chosen downstream question. Thus, the unrotated latent captures broad multimodal structure discovered by the model, whereas the rotated latent tightens that structure toward a specific analytical end point, such as stage, survival or ΔEF. The rotated dimensions should therefore be interpreted as endpoint-oriented views of a broader learnt signal, not as de novo discoveries independent of the chosen covariates.

The clinical variables chosen as rotation targets in each dataset and their encodings were as follows. For the IMMUcan NSCLC dataset, the clinical variables chosen as rotation targets were subtype, stage, sex, long survival, smoking history, and KRAS and TP53 mutation status. Long survival was encoded as a binary variable, with the threshold set to the approximate 80th percentile of survival time among deceased patients. This binary label was used only to orient one axis of the latent representation toward survival-related variation, as the rotation procedure itself does not accommodate censoring. Accordingly, the resulting survival-rotated latent should be interpreted as a survival-oriented analytical summary of a broader multimodal signal, rather than as a censored survival model. Downstream Cox and Kaplan–Meier analyses were then used to assess whether this survival-oriented summary was associated with patient outcome in a censoring-aware setting.

For consistency, we used the same rotation targets for the TCGA dataset, except for the mutation-status variables. For the ACS/CCS dataset, we used ΔEF, sex, classification (ACS, CCS, no CCS), and age. Because ACS patients contribute multiple timepoint-specific latent representations but only one measured EF value, the outcome-oriented rotation in this cohort should likewise be interpreted as a way of organising a broader recovered signal toward a clinically relevant end point, rather than as a direct longitudinal outcome model.

For the ACS/CCS dataset we used the Δ*EF* value; sex, male (1) vs. female (0); classification (ACS, CCS, no CCS); and patient age. The rotation for this dataset is not without noise, as ACS patients have multiple timepoint representations in the latent space but only one measured Δ*EF* value. This dataset design aspect also makes subsequent analyses more difficult, as these multiple timepoint representations have to map to the same outcome.

#### Signed *R*^2^

Since analysis of the loading matrices *W* for each latent factor modality pair by the loading magnitude will vary heavily based on predictor scales, and can be noisy for rotated factors, we introduce a signed *R*^2^ metric:

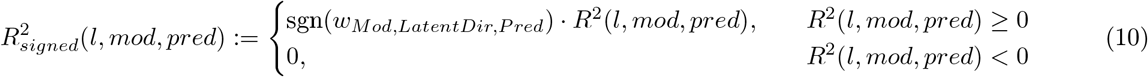

It is scale invariant, takes values in [−1, 1], where −1 indicates a perfectly reconstructed predictor with negative loading, 0 indicates a predictor reconstructed by mean values or worse, and 1 indicates a perfectly reconstructed predictor with positive loading. Importantly, high absolute values of 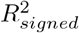 indicate a predictor that is well aligned with the loadings of the latent direction being investigated and was not just assigned a high loading value.

#### Statistical tests

We used sci-kit learn implementations of Analysis-of-variance (ANOVA) test, logistic regression, and linear regression. For Kaplan-Meier and Cox model estimates, we used lifelines implementation of KaplanMeierFitter and CoxPH-Fitter.

For multiple hypotheses testing, we used Benjamini-Hochberg (B-H) correction. Throughout the text, we refer to B-H corrected *p*-values as *q*-values.

#### GSEA

We used the GSEA implementation from fgsea R package [40] on the Reactome [22] C2 pathways, setting minSize = 15, maxSize = 500 and other parameters kept default.

#### Cluster celltype annotation

Cell clusters in ACS/CCS analysis were first annotated using positive marker genes (Supp. File 3) and then labels were refined by performing differential gene expression analysis between clusters and analysing the top 10 and bottom 10 genes for each cluster.

#### Simulation evaluation metrics

To evaluate the models performance on simulated data we used the following metrics.

For simple modalities, we defined modality specific *R*^2^ as

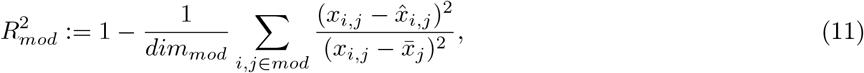

where *x*_*i,k*_ is the (pre-noise) *k*-th feature of modality *mod* for patient *i*, 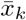 is its average over patients, and 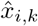 is its decoded prediction from the model.

Since the inferred clustering in structured modalities does not necessarily map one-to-one with the generative clustering, for structured modalities we defined modality specific *R*^2^ as

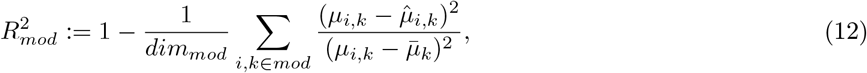

where *µ*_*i,k*_ are the *a posteriori* proportions of cluster *k* of modality *mod* for patient *i*, 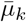 is its average over patients, and 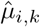 is its decoded prediction from the model. This formulation measures how much information from the inferred clustering structure proportions is included in the latent space of the model.

Then, we defined the total *R*^2^ as the weighted average of modality specific *R*^2^

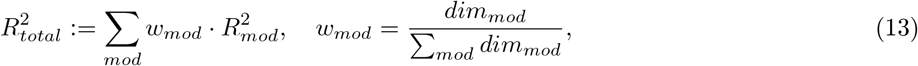

and mean view *R*^2^ as

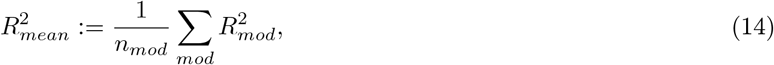

where *n*_*mod*_ is the total number of modalities. The first score measures how much total data signal was reconstructed in the model’s latent space, and the second measures how well all views were integrated in that latent.

For both the topic and GMM structured views, we used Adjusted Rand Index (ARI) between the labels from the generated clustering and the reconstructed clustering. Additionally, we defined clustering profiles distance score as:

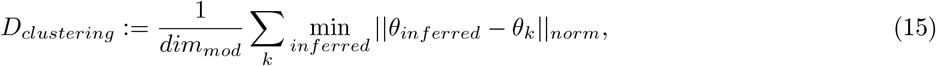

where *norm* is the modality appropriate *p*-norm, 2 for GMM and for topic modalities.

For the GMM modality, we used two separate clustering evaluation metrics, relative Calinski-Harabasz index (rel. CH) and relative silhouette score (rel. SS), both defined as the respective score of the inferred clustering divided by that score for the generative clustering.

For the topic modalities we used a separate clustering metric, relative loglikelihood (rel. LL), defined as

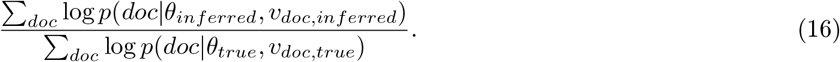

## Supporting information

Supplementary File 1 IMMUCAN patient modality availability

Supplementary File 2 Coronary syndrome patient modality availability

Supplemetary File 3 Positive celltype marker genes

## Data availability

IMMUcan NSCLC2 dataset will be available at [6]. TCGA LUAD and LUSC datasets are available at NCI Genomic Data Commons [33]. Coronary syndrome data is available at [5].

## Code availability

FACTMx model code is available at github.com/szczurek-lab/FACTMx.

## Author information

Conceptualization, K.O-O. and E.Sz.; Methodology, K.O-O. and E.Sz; Software, K.O-O.; Formal Analysis, K.O-O.; Investigation, K.O.O., M.Ł., Ł.K., D.W., M.Moż., (computational) and D.S., F.M., S.T., B.B.(experimental); Data Curation, D.S., F.M., M.Mou., S.T., B.B.; Writing – Original Draft, K.O-O. and E.Sz.; Writing – Review & Editing, all authors; Supervision, E.Sz.; Project Administration, H. S. H. and E.St.; Funding Acquisition, E.St. and E.Sz; Resources, R.L. and E.Sz.

All authors have read, edited and approved submission of the manuscript.

## Ethics declaration

Projects at Szczurek lab are co-founded by Merck Healthcare. H. S. H. and E.St. are full-time employees of the healthcare organization of Merck KGaA, Darmstadt, Germany. The remaining authors declare no competing interests.

## Supplementary Methods

### The FACTMx model

FACTMx is a variational autoencoder combining a flexible number of simple or structured heads modelling chosen data modalities.

Formally, let *z*_*i*_ ∈ ℝ^*L*^ be an *L*-dimensional latent representation vector for sample *i*, with an a priori distribution of *N* (0, *I*_*L*_). Let *s, k* be indices of simple and structured views respectively. We assume each sample *i* has observations *x*_*i,s*_ and *x*_*i,k*_ that are governed by the distributions of their heads. Let *v*_*k*_ be the collection of hidden variables of structured head *k* and *θ*_*k*_ be the directly optimisable parameters of that head.

For notation brevity we omit indices in cases where we want to indicate a set over the omitted index. For example, *z* would indicate the set of latent representations of all samples.

#### Simple heads

Simple heads model data that can be described with one distribution, with parameters that can be deterministically decoded from the latent representation of a sample or be optimized directly. Examples of simple heads are:

- Multinomial, such that *p*(*x*_*i,s*_|*z*_*i*_) ~ Multinomial(*ϕ*_*i,s*_); *ϕ*_*i,s*_ = *f*_*s*_(*z*_*i*_), where *x*_*i,s*_ is a vector of counts of category observations with probabilities *ϕ*_*i,s*_ that are decoded from *z*_*i*_ using a parametrisable function *f*_*s*_.
- Multinormal, such that *p*(*x*_*i,s*_|*z*_*i*_) ~ *N* (*µ*_*i,s*_, Σ_*i,s*_); (*µ*_*i,s*_, Σ_*i,s*_) = *f*_*s*_(*z*_*i*_) where *x*_*i,s*_ is a real number vector, *µ* is the mean, and Σ is the covariance matrix decoded from *z*_*i*_ using a parametrisable function *f*_*s*_.
- Poisson, such that *p*(*x*_*i,s*_ | *z*_*i*_) ~ Poisson(*µ*_*i,s*_); *µ*_*i,s*_ = *f*_*s*_(*z*_*i*_), where *µ*_*i,s*_ is the mean decoded from *z*_*i*_ using a parametrisable function *f*_*s*_.
- Zero Inflated Negative Binomial, such that *p*(*x*_*i,s*_ | *z*_*i*_) ~ ZINB(*ω*_*s*_, *ϕ*_*i,s*_, *r*_*i,s*_); (*ϕ*_*i,s*_, *r*_*i,s*_) = *f*_*s*_(*z*_*i*_), where *ω*_*s*_ is the zero inflation factor, optimised directly, *ϕ*_*i,s*_ are the success observation probabilities and *r*_*i,s*_ are the total trials, both decoded from *z*_*i*_ using a parametrisable function *f*_*s*_.

#### Structured heads

Structured heads model data that cannot be easily expressed with one tractable distribution but can be factorised into a probabilistic graphical model (PGM). For example, a structured head can be a mixture model, in which proportions of mixture components for a sample are decoded from its latent representation, and other parameters are optimised directly; in this publication we focus on these types of structured heads. As such, let us additionally define *y*_*i,k,d*_ as subobservation indexed with *d* that belongs to the data collection *x*_*i,k*_ of the structured view *k* for patient *i*; *χ*_*i,k,d*_ as the variable assigning this subobservation to one of the components of the structured view *k*; and *µ*_*i,k*_ as the decoded component proportions in view *k* for patient *i*. Then, some examples of the mixture-type structured heads are:

- Topic model, such that *x*_*i,k*_ is a collection of documents *d*, represented as vectors of word counts *y*_*i,k,d*_, that can belong to one of a set number of topics *t*, which are represented with word frequency profiles *β*_*k,t*_, such that

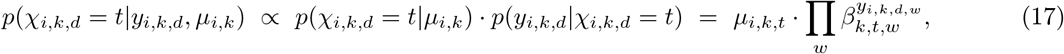

where *χ*_*i,k,d*_ is the assignment variable for document *d* in view *k* of sample *i, β*_*k,t,w*_ is the frequency of word *w* in topic *t* of view *k, y*_*i,k,d,w*_ is the count of word *w* in document *d* in view *k* of sample *i*, and *µ*_*i,k*_ is the a priori proportion of documents from topic *t* in view *k* of sample *i*, such that:

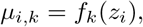

where *f*_*k*_ is a parametrisable function decoding *µ*_*i,k*_ from *z*_*i*_.
- ZINB mixture, such that *x*_*i,k*_ is a collection of subobservations *d*, represented as vectors of successful observation counts *y*_*i,k,d*_ that can belong to one of a set number of components *t*, which are parametrised by *ϕ*_*k,t,w*_, *r*_*k,t,w*_, *ω*_*k,t,w*_, where *ϕ*_*k,t,w*_ are the success probabilities at position *w* of view *k* in component *t, r*_*k,t,w*_ is the ratio of trials at position *w* of view *k* in component *t*, and *ω*_*k,t,w*_ is the inflation factor at position *w* of view *k* in component *t*, optimised directly. For the subobservation assignment probabilities we have:

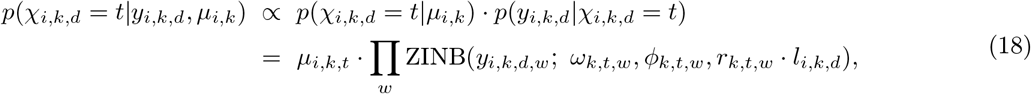

where *l*_*i,k,d*_ is the size factor for subobservation *d* of view *k* of sample *i, r*_*k,t,w*_ · *l*_*i,k,t*_ is the total number of trials at position *w* in component *t* for subobservation *d* of view *k* of sample *i*, and *µ*_*i,k*_ is the a priori proportion of subobservations from component *t* in view *k* of sample *i*, such that:

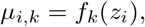

where *f*_*k*_ is a parametrisable function decoding *µ*_*i,k*_ from *z*_*i*_.
- mixture of Gaussians, such that *x*_*i,k*_ is a collection of subobservations *d*, represented as real number vectors *y*_*i,k,d*_, that can belong to one of a set number of components *t*, which are parametrised by *l*_*k,t*_, Σ_*k,t*_, where *l*_*k,t*_ is the location of component *t* in view *k*, Σ_*k,t*_ is the covariance matrix for component *t* in view *k*. For the subobservation assignment probabilities we have:

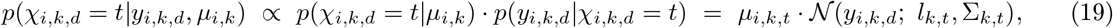

where *µ*_*i,k*_ is the a priori proportion of subobservations from component *t* in view *k* of sample *i*, such that:

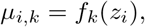

where *f*_*k*_ is a parametrisable function decoding *µ*_*i,k*_ from *z*_*i*_.

Due to flexibility of the variational inference, the parameter decoding functions *f*_*k*_ can be trainable MLPs or any deep learning models. If the available data is scarce on the observation level (*i*) we can use a one layer MLP with no activation, which imposes linear dependency of the parameters on the latent factors, similarly to a MOFA model.

#### General graph formulation

In general, every FACTMx model can be represented as an encoder-decoder pair of graphs.

The decoder graph models *p*, the generative distribution underlying the data. The complete distribution of *p* for sample *i* factorises as:

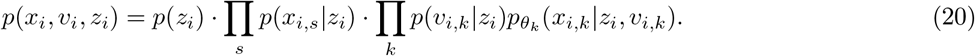

If only mixture-type structured heads are used, this becomes:

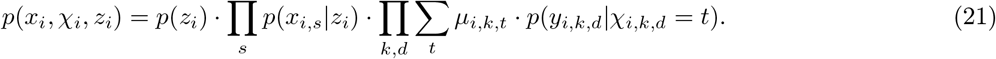

Since the posterior distribution *p*(*z*_*i*_, *v*_*i*_|*x*_*i*_) is intractable, we approximate it with a variational distribution *q*, modelled with the encoder graph. It is formulated so that its complete distribution factorises as:

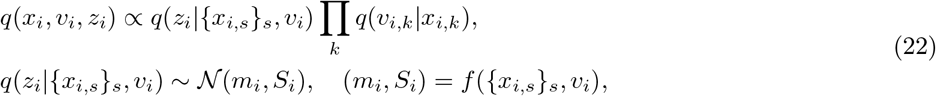

where *m*_*i*_ is the mean, and *S*_*i*_ is the covariance matrix decoded from ({*x*_*i,s*_}_*s*_, *v*_*i*_) using a parametrisable function *f*. To obtain a model *q* with this factorisation it is sufficient that no direct dependencies exist between heads and none of variables from *v* are children of *z*.

With only mixture models as structured heads, we can reformulate eq. 22 as:

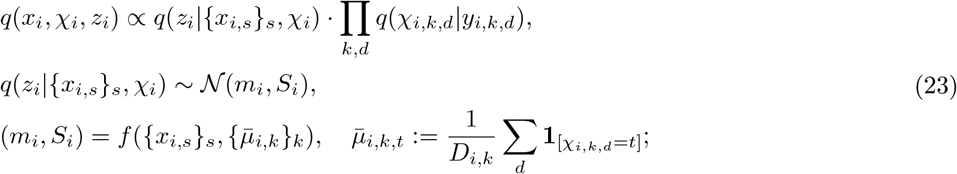

where *D*_*i,k*_ is the number of subobservations in structured view *k* for sample *i*. The variational assignment distribution *q*(*χ*_*i,k,d*_ = *t*|*y*_*i,k,d*_) can be defined in any way; we use the most practical approach, which is a deep neural classifier that takes as input the subobservation data and outputs the classification logits.

To infer the parameters of *p* and *q* we use the evidence lower bound (ELBO, ℒ) objective. Due to conditional independence of heads in *p* given *z* and the form of *q* in eq. 22, the ELBO for sample *i* factorises as:

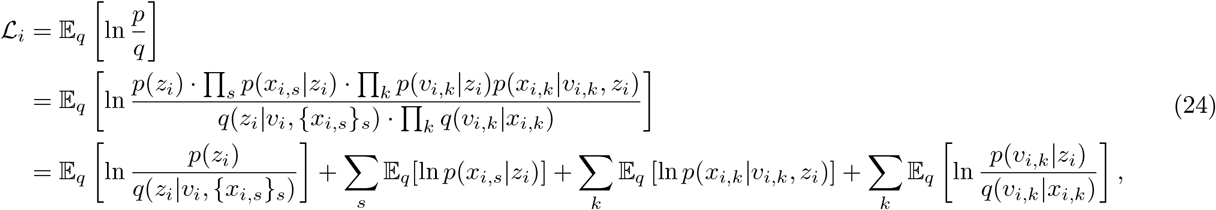

where the total ELBO is ℒ= ∑_*i*_ ℒ_*i*_. Note that the first term is equivalent to − KL(*q*(*z*_*i*_|*v*_*i*_, {*x*_*i,s*_}_*s*_) ∥ *p*(*z*_*i*_)), the KL divergence between the variational posterior and the prior over *z*_*i*_. Similarly, if variables in *v*_*i,k*_ form a Bayesian network in both *p* and *q*, then the last term can be decomposed as:

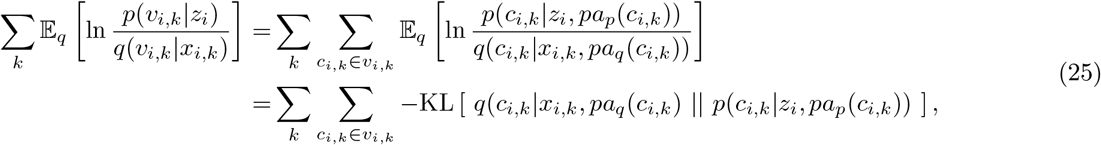

where *c*_*i,k*_ are the variables forming the set *v*_*i,k*_ and *pa*_*p*_ and *pa*_*q*_ denote parent variables in *p* and *q* respectively.

Again, with only mixture models we can further derive this formulation:

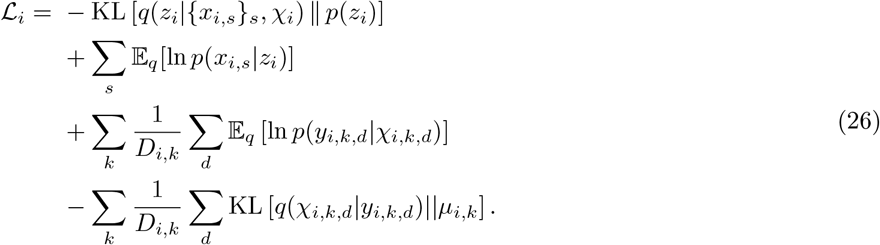

Note that last term denotes the KL divergence between the variational assignment distribution and the decoded per-sample proportions.

Gradients of ℒ are approximated through the VAE gradient procedure (assuming reparametrisation trick) using algorithm 1 in a general setting and algorithm 2 in a setting with only mixture-type structured heads.

##### Algorithm 1

Stochastic Gradient Estimation of ℒ

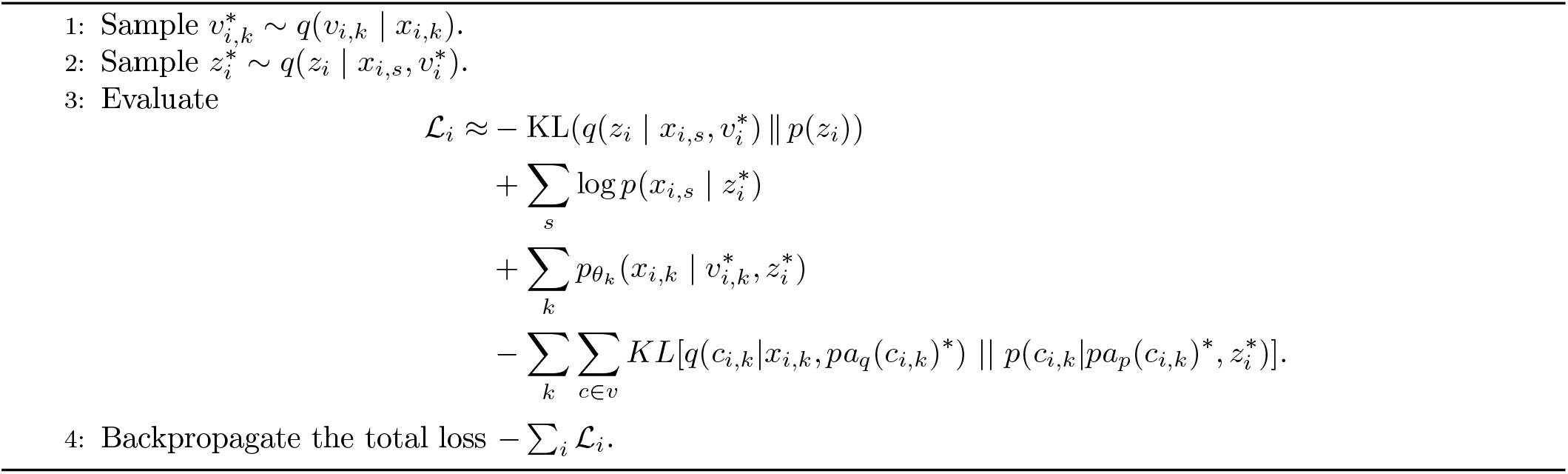

##### Algorithm 2

Stochastic Gradient Estimation of ℒ with mixture-type structured heads

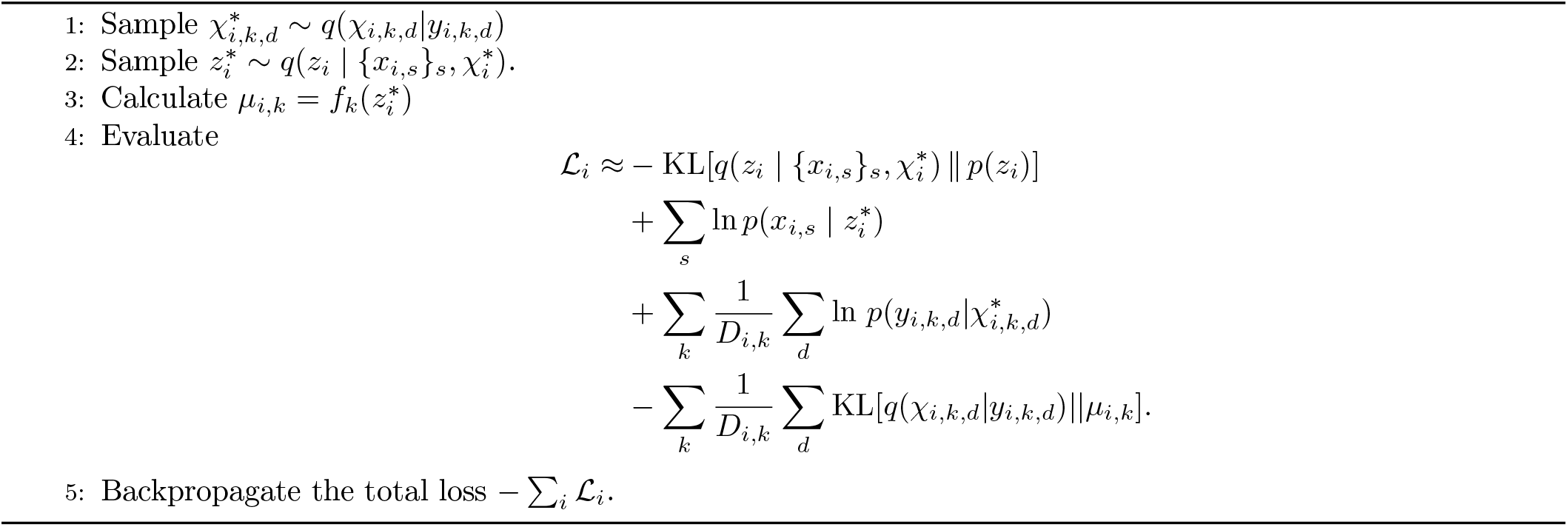

### Data simulation

#### Simulation parameters

The simulation scenarios are governed by two parameters.

The first is noise scale 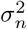, with values adjusted such that the *R*^2^ of the resulting simple data is ≈ 1 for 0% noise scenarios, ≈ 0.8 for 20% noise scenarios, and ≈ 0.5 for 50% noise scenarios.

The second is the sparsity of the dependency matrices. If sparsity is used, then weights of the dependency matrices *W*_*mod*_ are zeroed out for columns mapping to specific modality-factor combinations, which means those factors no longer impact the data of those modalities.

We use six simulation scenarios resulting from the combination of noise levels and sparsity type.

#### Simulation process

Assume we are given:

- number of patients *N*_*patients*_,
- number of observations for structured views *N*_*subobs*_,
- number of true latent factors *dim*_*latent*_,
- for each modality *mod* its dimensionality *dim*_*mod*_ (for the structured modalities this is the number of clusters),
- for each structured modality the dimensionality of its observations *dim*_*obs,mod*_.

Then the simulation setup is:

- Step 1: simulate dependency matrices *W*_*mod*_:
  1. Draw weights for each dependency from 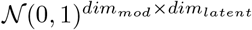
  2. For sparse dependency scenarios, for each modality zero out the weights in the columns of *W*_*mod*_ that correspond to unrelated factors
- Step 2: simulate additional parameters of structured modalities
  1. For topic modalities: draw topic profiles *θ* for each topic modality *mod* using algorithm 3
  2. For GMM modality *mod*:
    – initialise the centers *l*_*mod*_ of the GMM mixture components using an orthogonal matrix obtained from the QR decomposition of a matrix of random numbers drawn from a normal distribution,
    – scale the centers *l*_*mod*_ by 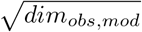,
    – initialise the covariance matrix Σ_*mod,c*_ for each component of the mixture with diagonal entries such that Σ_*mod,c,i,i*_ = |*l*_*mod,c,i*_| + 0.5 and off-diagonal entries as 0.
- Step 3: Patient simulation
  1. Simulate latent representations *z*_*i*_ ~ *N* (0, Σ), such that Σ is a diagonal matrix with entries 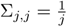. This results in latent directions that explain linearly less variance of the data.
  2. Obtain parameters of patient’s simple views from the latent using *W*_*mod*_ · *z*_*i*_
  3. Draw simple view data for patients from their respective data distributions, then add gaussian noise with the simulations noise parameter
  4. For each structured view *k*:
    – obtain the patient observation proportions using *µ*_*i,k*_ = Softmax(*W*_*k*_ · *z*_*i*_)
    – For observations from structured modalities, draw the observation assignments *ξ* from a categorical distribution with probabilities *µ*_*i,k*_
    – Generate the data *x*_*i,k*_

##### Algorithm 3

Topic Profile Generation per Modality

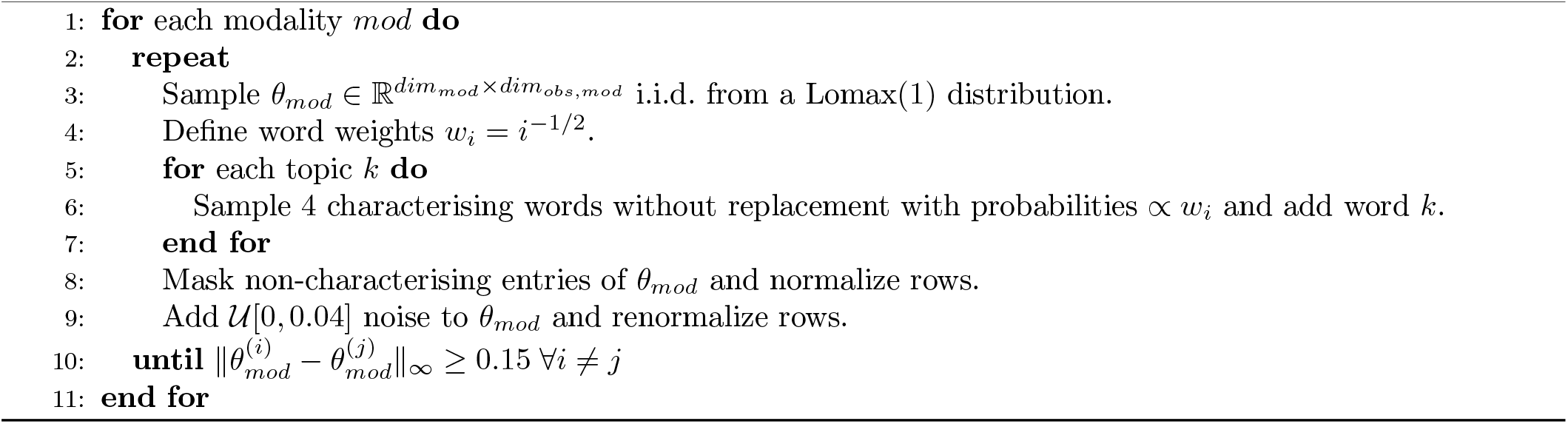

## References

1. Wilk, A. J., Rustagi, A., Zhao, N. Q., Roque, J., Martínez-Colón, G. J., McKechnie, J. L., Ivison, G. T., Ranganath, T., Vergara, R., Hollis, T., Simpson, L. J., Grant, P., Subramanian, A., Rogers, A. J. & Blish, C. A. A single-cell atlas of the peripheral immune response in patients with severe COVID-19. en. Nat. Med. 26, 1070–1076 (July 2020).

2. Keren, L., Bosse, M., Marquez, D., Angoshtari, R., Jain, S., Varma, S., Yang, S.-R., Kurian, A., Van Valen, D., West, R., Bendall, S. C. & Angelo, M. A structured tumor-immune microenvironment in triple negative breast cancer revealed by multiplexed ion beam imaging. en. Cell 174, 1373–1387.e19 (Sept. 2018).

3. Ayers, M., Lunceford, J., Nebozhyn, M., Murphy, E., Loboda, A., Kaufman, D. R., Albright, A., Cheng, J. D., Kang, S. P., Shankaran, V., Piha-Paul, S. A., Yearley, J., Seiwert, T. Y., Ribas, A. & McClanahan, T. K. IFN-γ-related mRNA profile predicts clinical response to PD-1 blockade. en. J. Clin. Invest. 127, 2930–2940 (Aug. 2017).

4. Ridker, P. M., Hennekens, C. H., Buring, J. E. & Rifai, N. C-reactive protein and other markers of inflammation in the prediction of cardiovascular disease in women. en. N. Engl. J. Med. 342, 836–843 (Mar. 2000).

5. Pekayvaz, K. et al. Multiomic analyses uncover immunological signatures in acute and chronic coronary syndromes. en. Nat. Med. 30, 1696–1710 (June 2024).

6. Schulz, D. et al. Comprehensive characterization of early-stage Non-Small Cell Lung cancers with multi-modal data integration Nov. 2025.

7. The Cancer Genome Atlas Research Network. Comprehensive genomic characterization of squamous cell lung cancers. Nature 489, 519–525 (2012).

8. The Cancer Genome Atlas Research Network. Comprehensive molecular profiling of lung adenocarcinoma. Nature 511, 543–550 (2014).

9. Ilse, M., Tomczak, J. M. & Welling, M. Attention-based deep multiple instance learning (Feb. 2018).

10. Chen, R. J., Lu, M. Y., Wang, J., Williamson, D. F. K., Rodig, S. J., Lindeman, N. I. & Mahmood, F. Pathomic Fusion: An integrated framework for fusing histopathology and genomic features for cancer diagnosis and prognosis. en. IEEE Trans. Med. Imaging 41, 757–770 (Apr. 2022).

11. Blei, D. M., Ng, A. Y. & Jordan, M. I. Latent Dirichlet Allocation. Journal of Machine Learning Research 3, 993–1022 (2003).

12. Traag, V. A., Waltman, L. & van Eck, N. J. From Louvain to Leiden: guaranteeing well-connected communities. en. Sci. Rep. 9, 5233 (Mar. 2019).

13. Argelaguet, R., Velten, B., Arnol, D., Dietrich, S., Zenz, T., Marioni, J. C., Buettner, F., Huber, W. & Stegle, O. Multi-Omics Factor Analysis-a framework for unsupervised integration of multi-omics data sets. en. Mol. Syst. Biol. 14, e8124 (June 2018).

14. Argelaguet, R., Arnol, D., Bredikhin, D., Deloro, Y., Velten, B., Marioni, J. C. & Stegle, O. MOFA+: a statistical framework for comprehensive integration of multi-modal single-cell data. en. Genome Biol. 21, 111 (May 2020).

15. Rodriguez, L. et al. Systems-Level Immunomonitoring from Acute to Recovery Phase of Severe COVID-19. Cell Reports Medicine 1, 100078 (2020).

16. Consiglio, C. R. et al. The Immunology of Multisystem Inflammatory Syndrome in Children with COVID-19. Cell 183, 968–981.e7 (2020).

17. Lerma-Martin, C. et al. Cell Type Mapping Reveals Tissue Niches and Interactions in Subcortical Multiple Sclerosis Lesions. Nature Neuroscience 27, 2354–2365 (2024).

18. Łazęcka, M. & Szczurek, E. M. Factor Analysis with Correlated Topic Model for Multi-Modal Data in Proceedings of The 28th International Conference on Artificial Intelligence and Statistics (eds Li, Y., Mandt, S., Agrawal, S. & Khan, E.) 258 (PMLR, May 2025), 1801–1809. https://proceedings.mlr.press/v258/lazecka25a.html.

19. Chen, R. J., Ding, T., Lu, M. Y., Williamson, D. F. K., Jaume, G., Chen, B., Zhang, A., Shao, D., Song, H., Shaban, M., et al. Towards a General-Purpose Foundation Model for Computational Pathology. Nature Medicine (2024).

20. HUGO Gene Nomenclature Committee. HGNC Database RRID:SCR002827. https://www.genenames.org (2026).

21. Subramanian, A., Tamayo, P., Mootha, V. K., Mukherjee, S., Ebert, B. L., Gillette, M. A., Paulovich, A., Pomeroy, S. L., Golub, T. R., Lander, E. S. & Mesirov, J. P. Gene set enrichment analysis: A knowledge-based approach for interpreting genome-wide expression profiles. Proceedings of the National Academy of Sciences of the United States of America 102, 15545–15550 (2005).

22. Reactome. Reactome Pathway Database https://reactome.org (2025).

23. Wu, J. & Chen, Y. Unraveling the Connection: Extracellular Vesicles and Non-Small Cell Lung Cancer. International Journal of Nanomedicine 19, 8139–8157 (2024).

24. Diao, X., Guo, C., Zheng, H., Zhao, K., Luo, Y., An, M., et al. SUMOylation-triggered ALIX activation modulates extracellular vesicles circTLCD4-RWDD3 to promote lymphatic metastasis of non-small cell lung cancer. Signal Transduction and Targeted Therapy 8, 426 (2023).

25. Qian, C., Jiang, Z., Zhou, T., Wu, T., Zhang, Y., Huang, J., Ouyang, J., Dong, Z., Wu, G. & Cao, J. Vesiclemediated transport-related genes are prognostic predictors and are associated with tumor immunity in lung adenocarcinoma. Frontiers in Immunology 13, 1034992 (2022).

26. House, I. G., Savas, P., Lai, J., Chen, A. X. Y., Oliver, A. J., Teo, Z. L., et al. Macrophage-Derived CXCL9 and CXCL10 Are Required for Antitumor Immune Responses Following Immune Checkpoint Blockade. Clinical Cancer Research 26, 487–504 (2020).

27. Bill, R., Wirapati, P., Messemaker, M., Roh, W., Zitti, B., Duval, F., et al. CXCL9:SPP1 macrophage polarity identifies a network of cellular programs that control human cancers. Science 381, 515–524 (2023).

28. Kuang, D.-M., Zhao, Q., Wu, Y., Peng, C., Wang, J., Xu, Z.Yin, X.-Y. & Zheng, L. Peritumoral neutrophils link inflammatory response to disease progression by fostering angiogenesis in hepatocellular carcinoma. Journal of Hepatology 54, 948–955 (2011).

29. Zhou, J., Xu, Q., Liu, H., Miao, J., Bian, C., Wei, Y., Wang, W. & Jiang, S. Prognostic value of tumor-associated CD177+ neutrophils in lung adenocarcinoma. Oncology Letters 27, 189 (2024).

30. Pedregosa, F. et al. Scikit-learn: Machine Learning in Python. Journal of Machine Learning Research 12. Accessed: 2026-05, 2825–2830. https://scikit-learn.org/stable/ (2011).

31. Eling, N. et al. Multi-modal image analysis for large-scale cancer tissue studies within IMMUcan. Cell Reports Methods 5. Accessed: 2026-05, 101170. https://www.cell.com/cell-reports-methods/fulltext/S2667-2375(25)00206-1 (mSept. 2025).

32. Zhang, A., Jaume, G., Vaidya, A., Ding, T. & Mahmood, F. Accelerating Data Processing and Benchmarking of AI Models for Pathology. arXiv preprint arXiv:2502.06750. TRIDENT toolkit/pipeline for large-scale whole-slide image processing; accessed: 2026-05. https://github.com/mahmoodlab/TRIDENT (2025).

33. Heath, A. P., Ferretti, V., Agrawal, S., et al. The NCI Genomic Data Commons. Nature Genetics 53. Accessed: 2026-05, 257–262 (2021).

34. NCI Genomic Data Commons. GDC Bioinformatics Pipeline: mRNA Analysis https://docs.gdc.cancer.gov/Data/Bioinformatics_Pipelines/Expression_mRNA_Pipeline/. Accessed: 2026-05. 2026.

35. NCI Genomic Data Commons. GDC Bioinformatics Pipeline: DNA-Seq Analysis: Whole Exome and Targeted Sequencing Variant Calling https://docs.gdc.cancer.gov/Data/Bioinformatics_Pipelines/DNA_Seq_Variant_Calling_Pipeline/. Accessed: 2026-05. 2026.

36. NCI Genomic Data Commons. GDC Bioinformatics Pipeline: Methylation Analysis https://docs.gdc.cancer.gov/Data/Bioinformatics_Pipelines/Methylation_Pipeline/. Accessed: 2026-05. 2026.

37. Hie, B., Bryson, B. & Berger, B. Efficient integration of heterogeneous single-cell transcriptomes using Scanorama. Nature Biotechnology 37, 685–691 (2019).

38. Kingma, D. P. & Ba, J. Adam: A Method for Stochastic Optimization. International Conference on Learning Representations. https://arxiv.org/abs/1412.6980 (2015).

39. Argelaguet, R., Velten, B., Arnol, D., Dietrich, S., Zenz, T., Marioni, J. C., Buettner, F., Huber, W. & Stegle, O. Multi-Omics Factor Analysis-a framework for unsupervised integration of multi-omics data sets. en. Mol. Syst. Biol. 14, e8124 (June 2018).

40. Bioconductor Package Maintainers. fgsea: Fast Gene Set Enrichment Analysis Bioconductor package. 2026.

