## Supplementary Figures for "Modelling interpretable patient-level representations from structured and simple multimodal data"

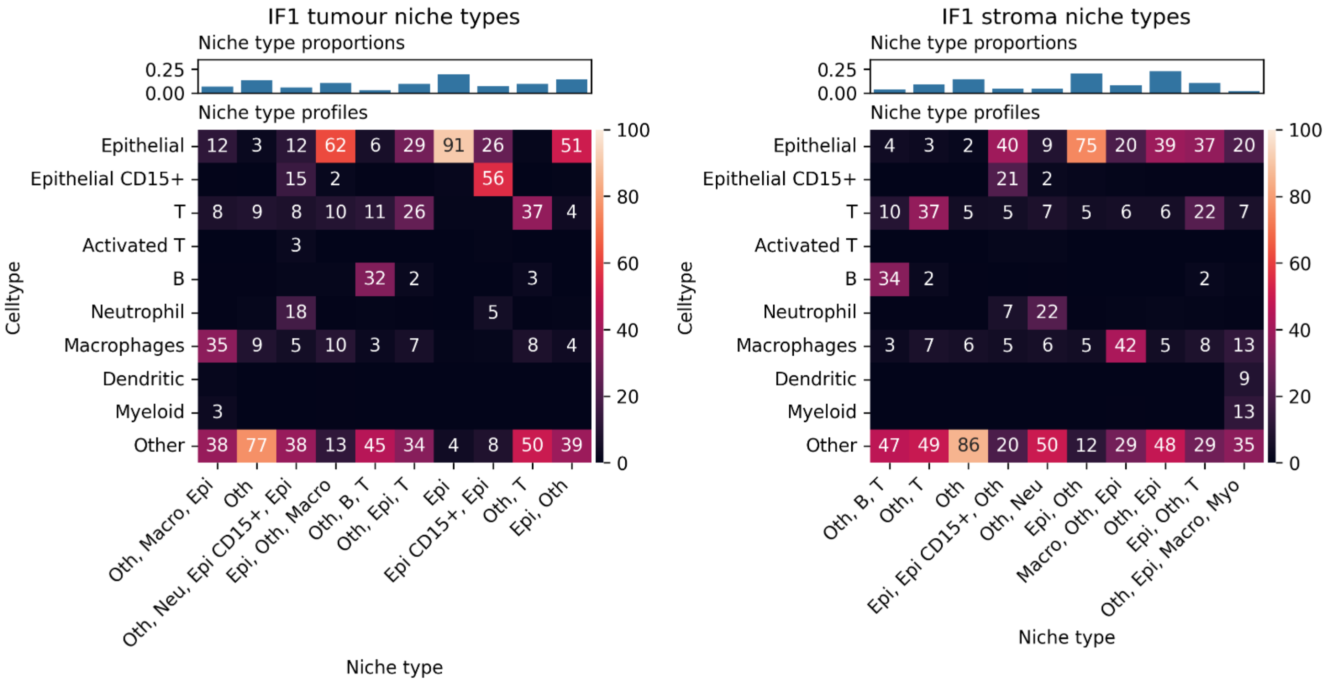


Supplementary Figure 1: Profiles of niche types in IF1 tumour and stroma regions

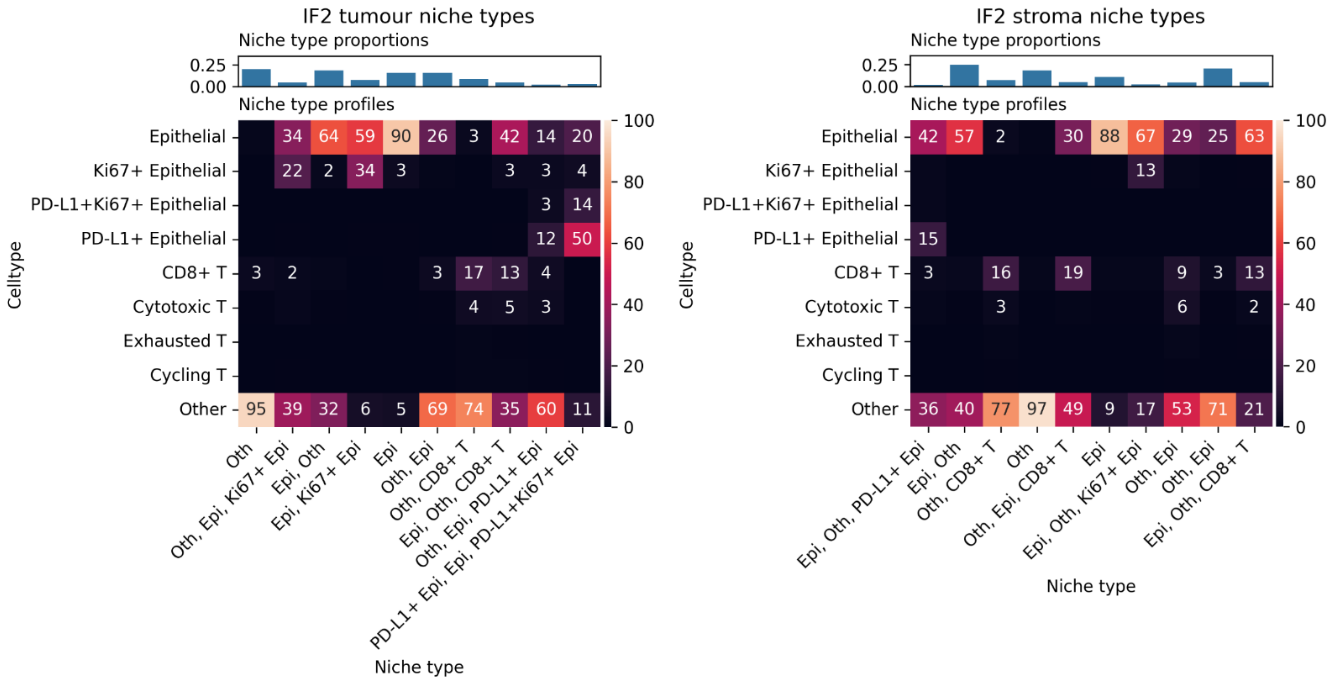


Supplementary Figure 2: Profiles of niche types in IF2 tumour and stroma regions

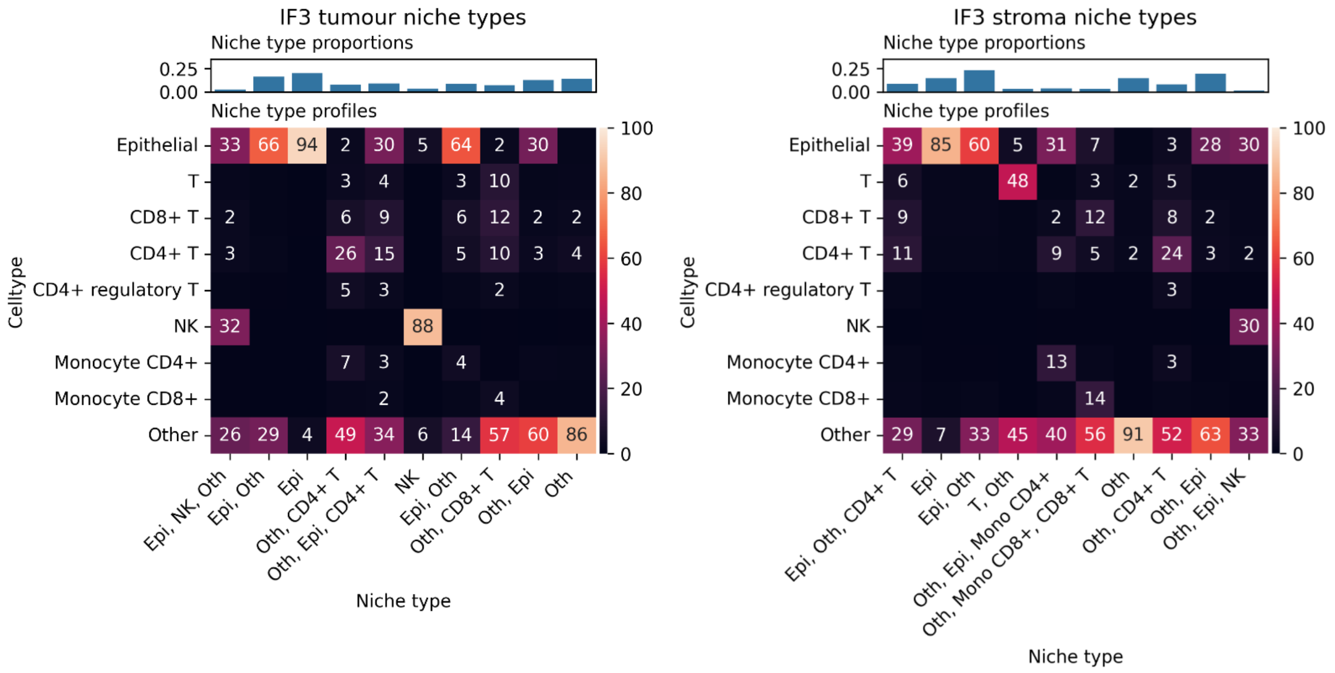


Supplementary Figure 3: Profiles of niche types in IF3 tumour and stroma regions

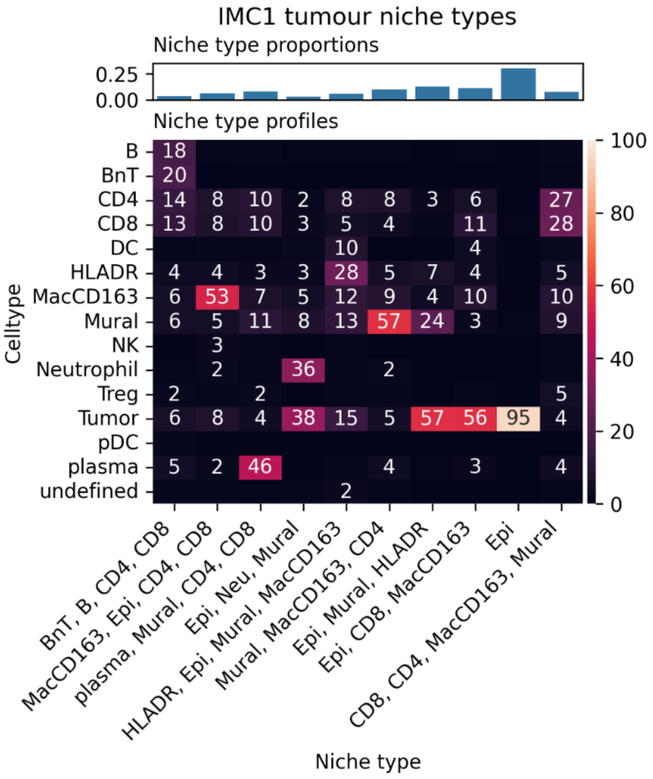

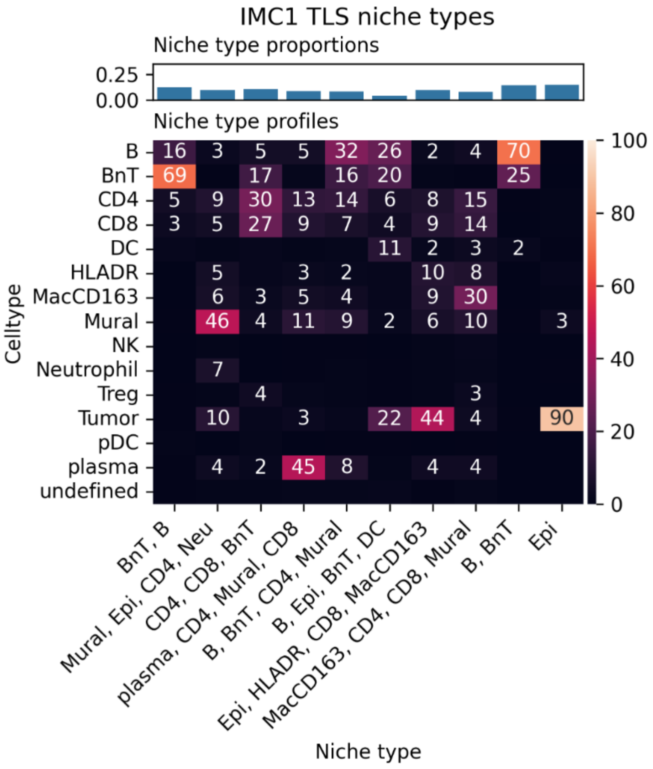

Supplementary Figure 4: Profiles of niche types in IMC1 tumour and TLS regions

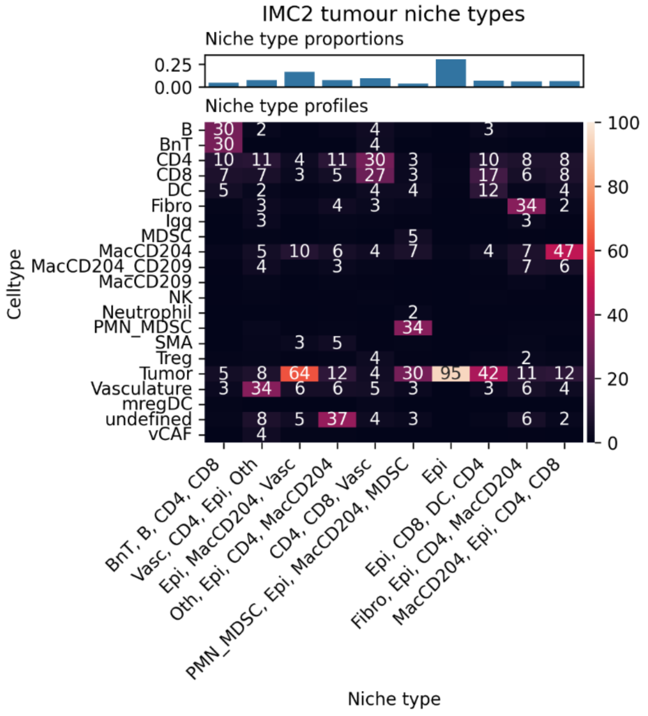

Supplementary Figure 5: Profiles of niche types in IMC2 tumour regions

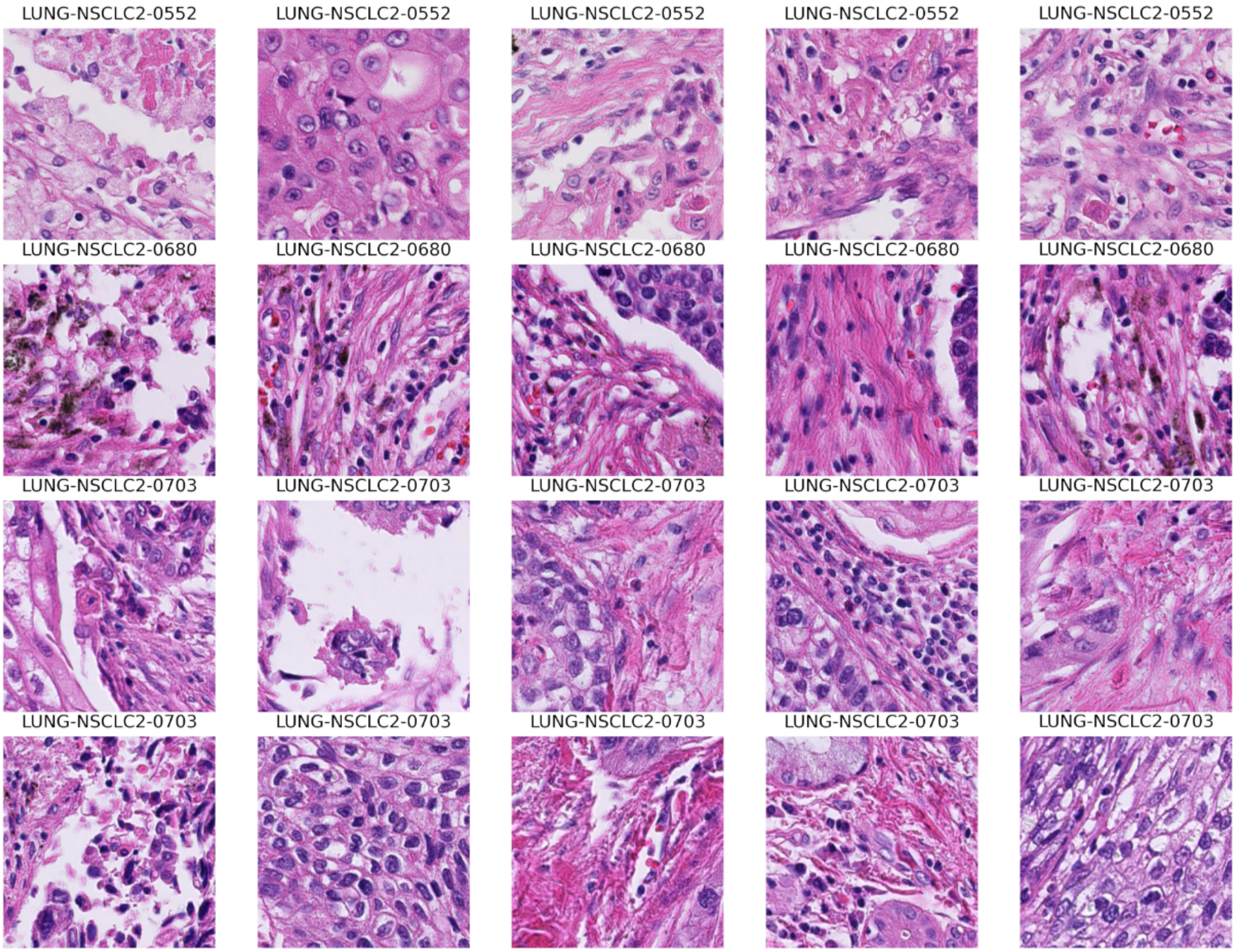


Supplementary Figure 6: Examples of H&E patches from cluster 0 – stroma-tumour border.

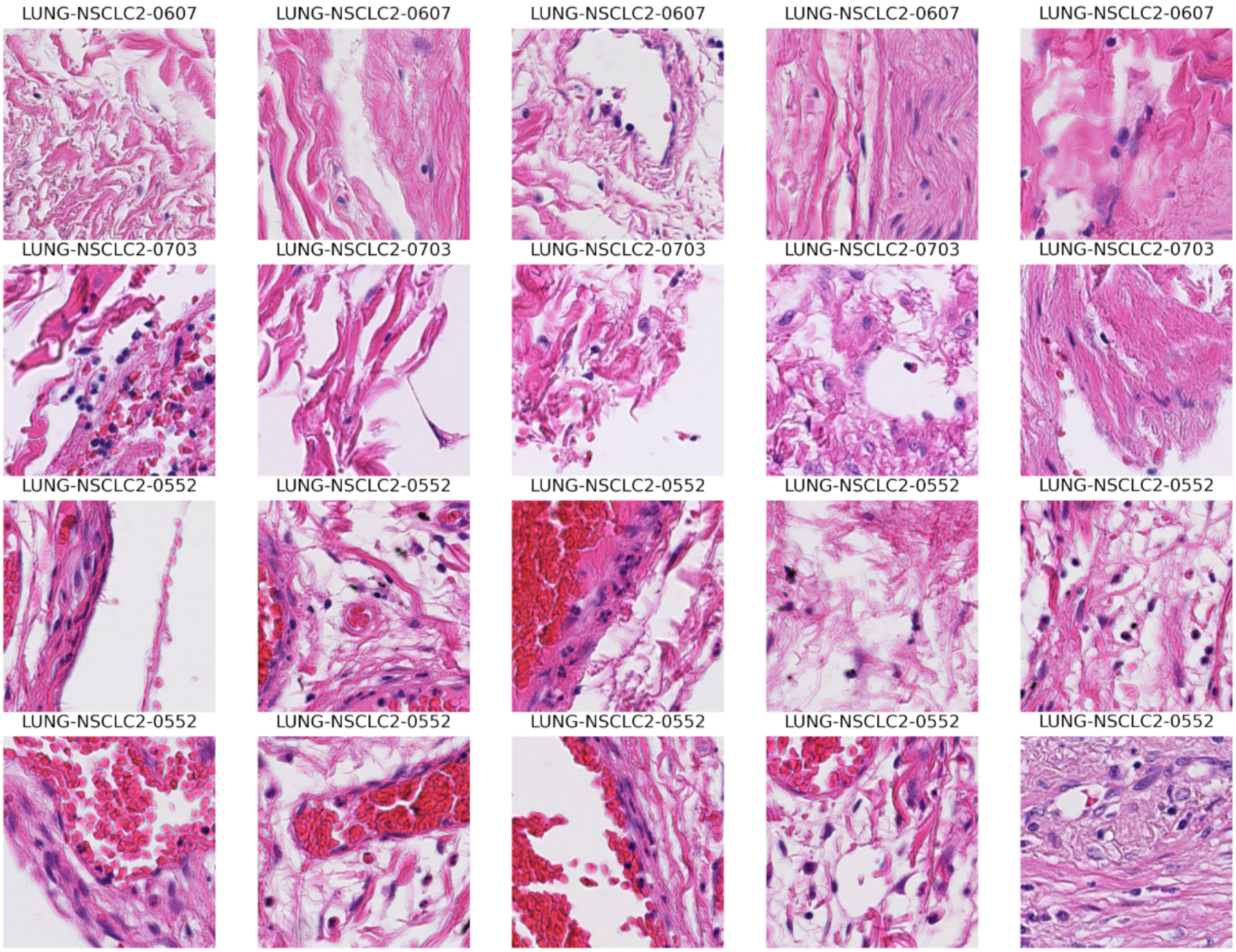


Supplementary Figure 7: Examples of H&E patches from cluster 1 – blood-rich peripheral tissue.

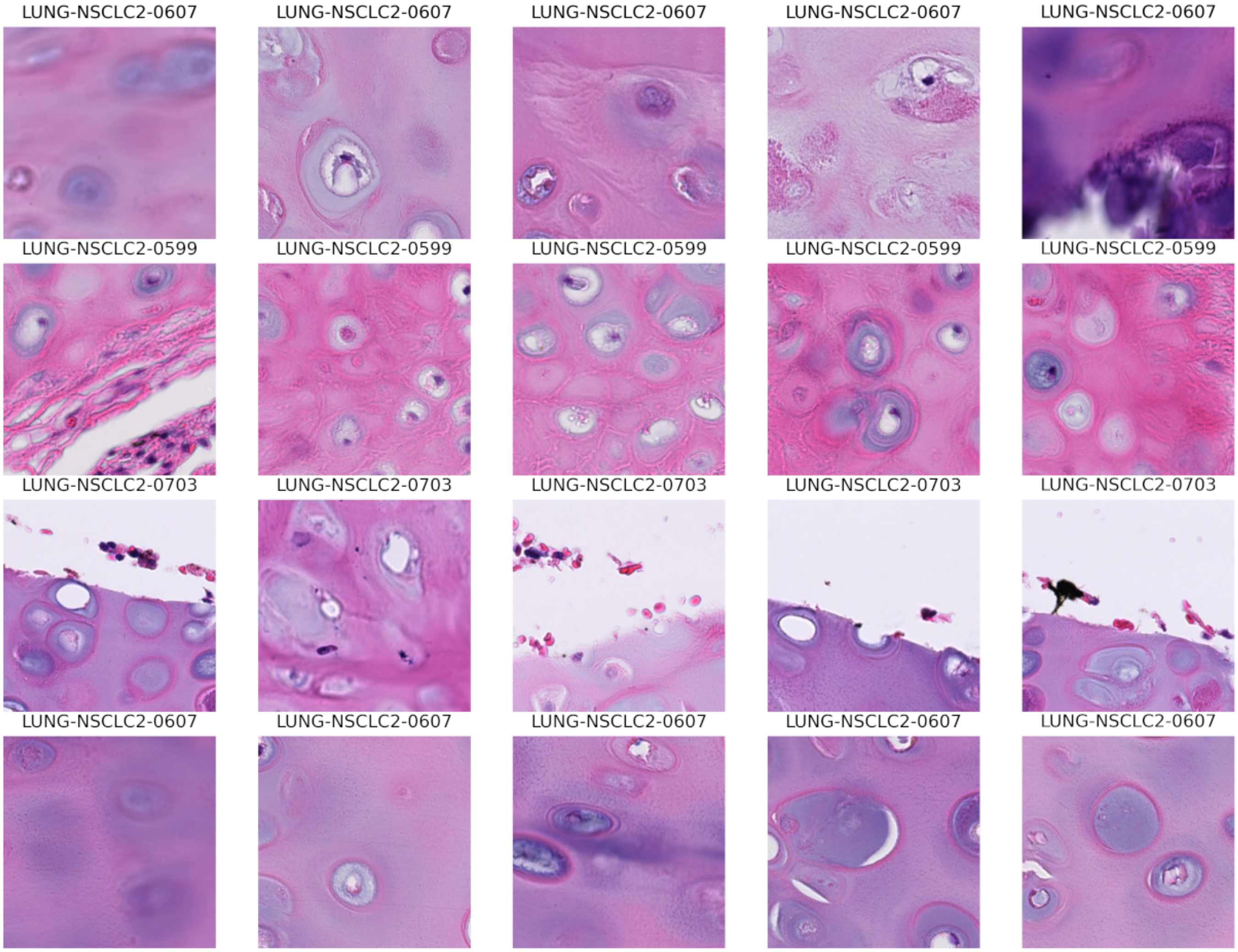


Supplementary Figure 8: Examples of H&E patches from cluster 2 – airway structural compartment.

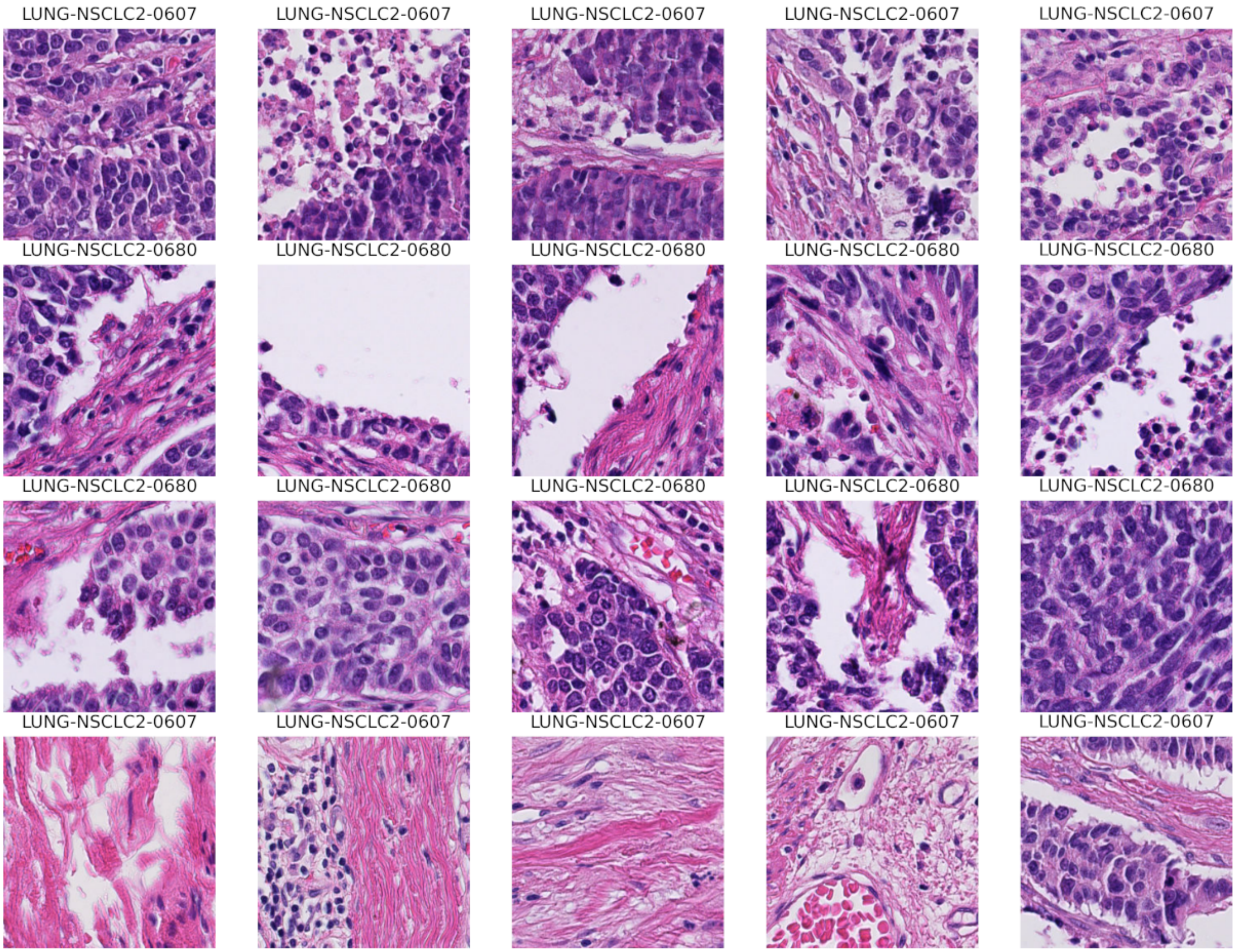


Supplementary Figure 9: Examples of H&E patches from cluster 3 – viable tumour epithelium.

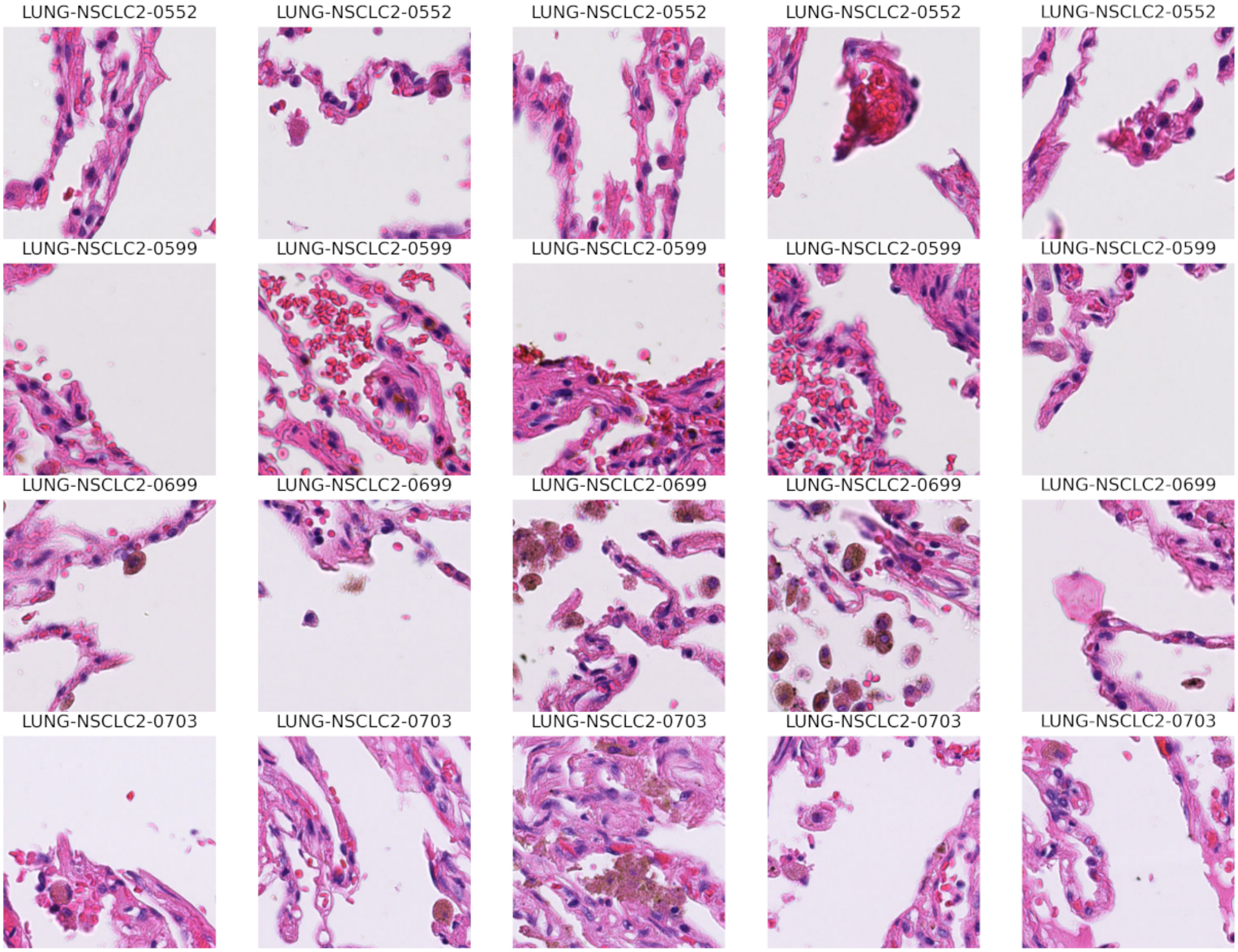


Supplementary Figure 10: Examples of H&E patches from cluster 4 – alveolar air-space.

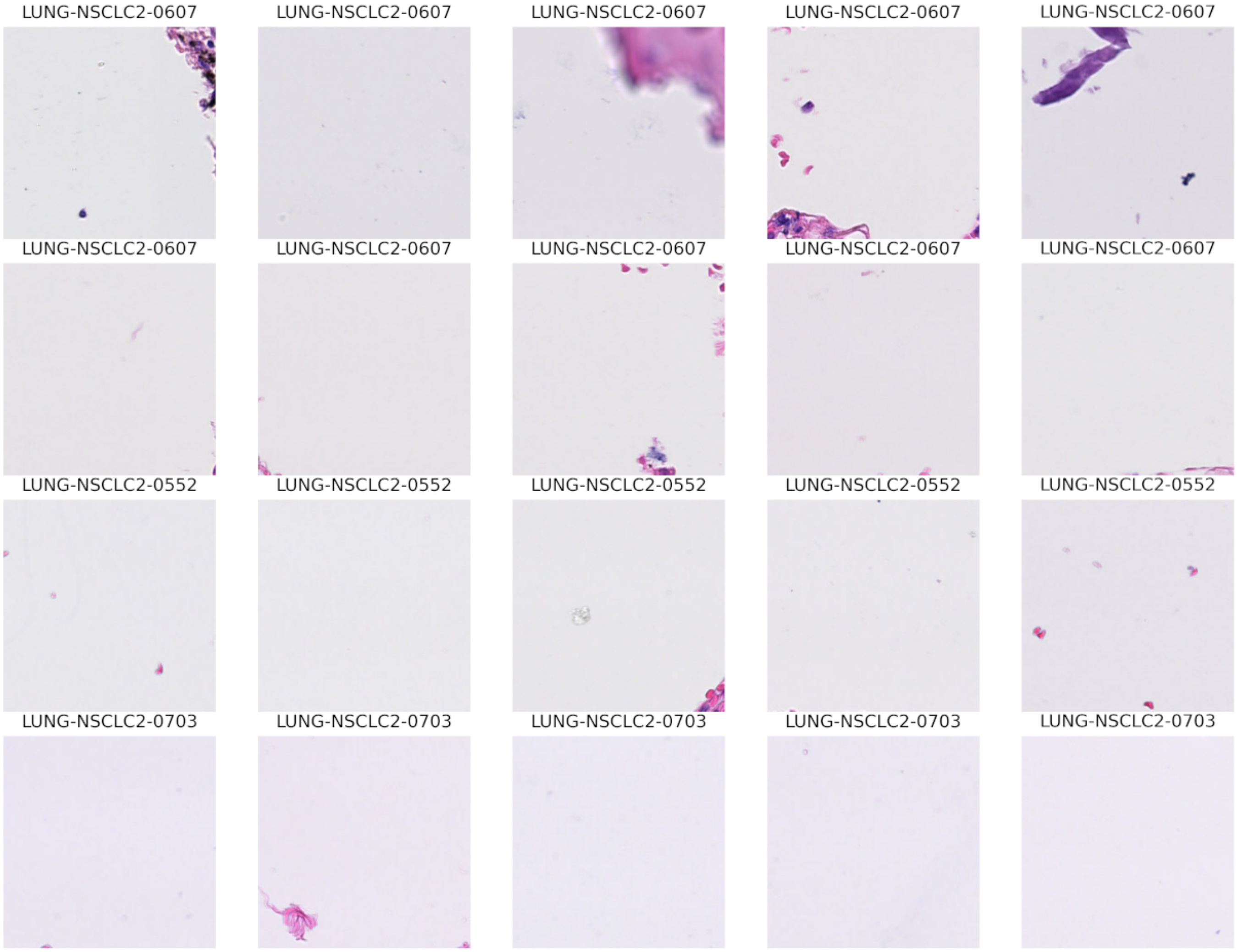


Supplementary Figure 11: Examples of H&E patches from cluster 5 – tissue edge.

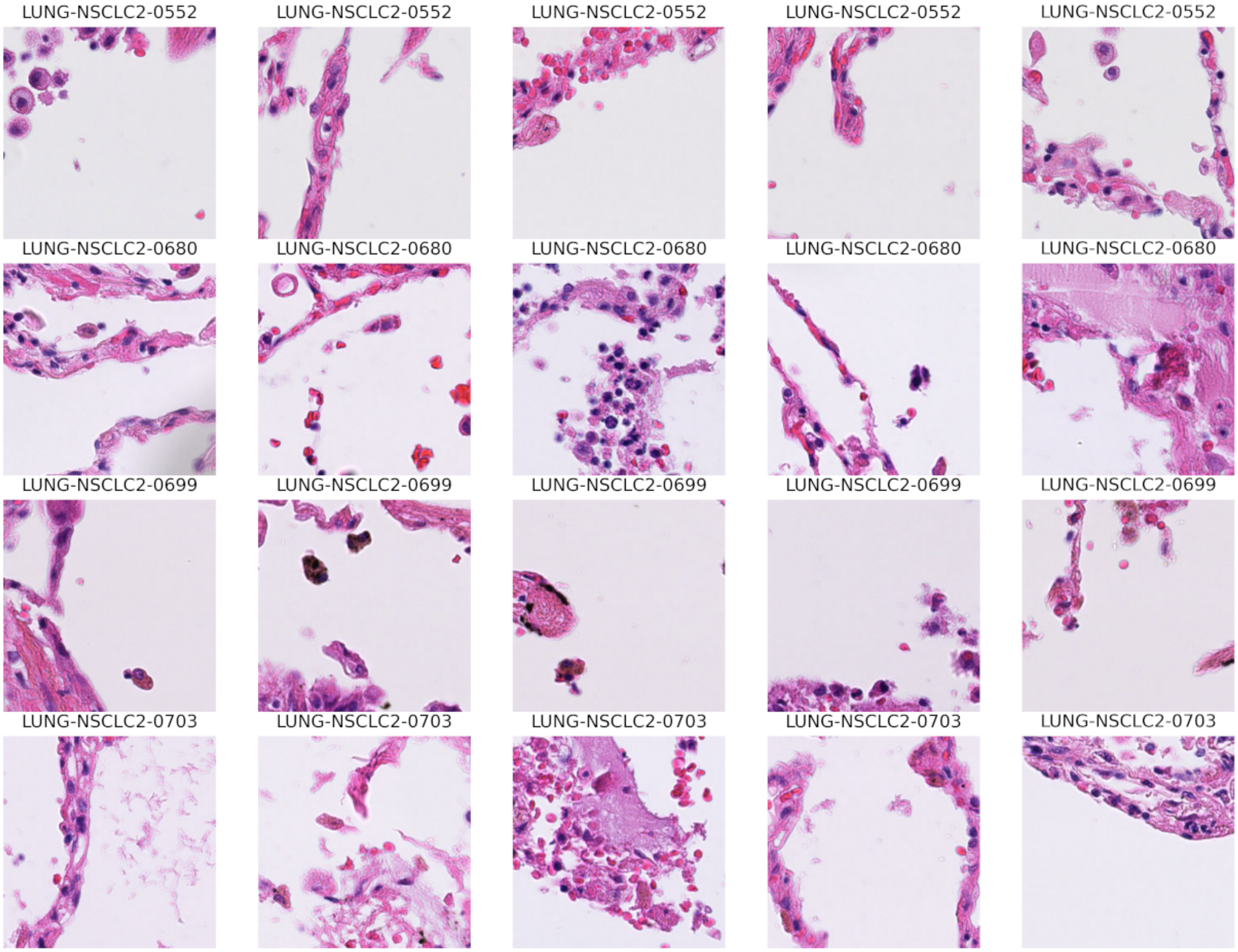


Supplementary Figure 12: Examples of H&E patches from cluster 6 – adjacent normal lung.

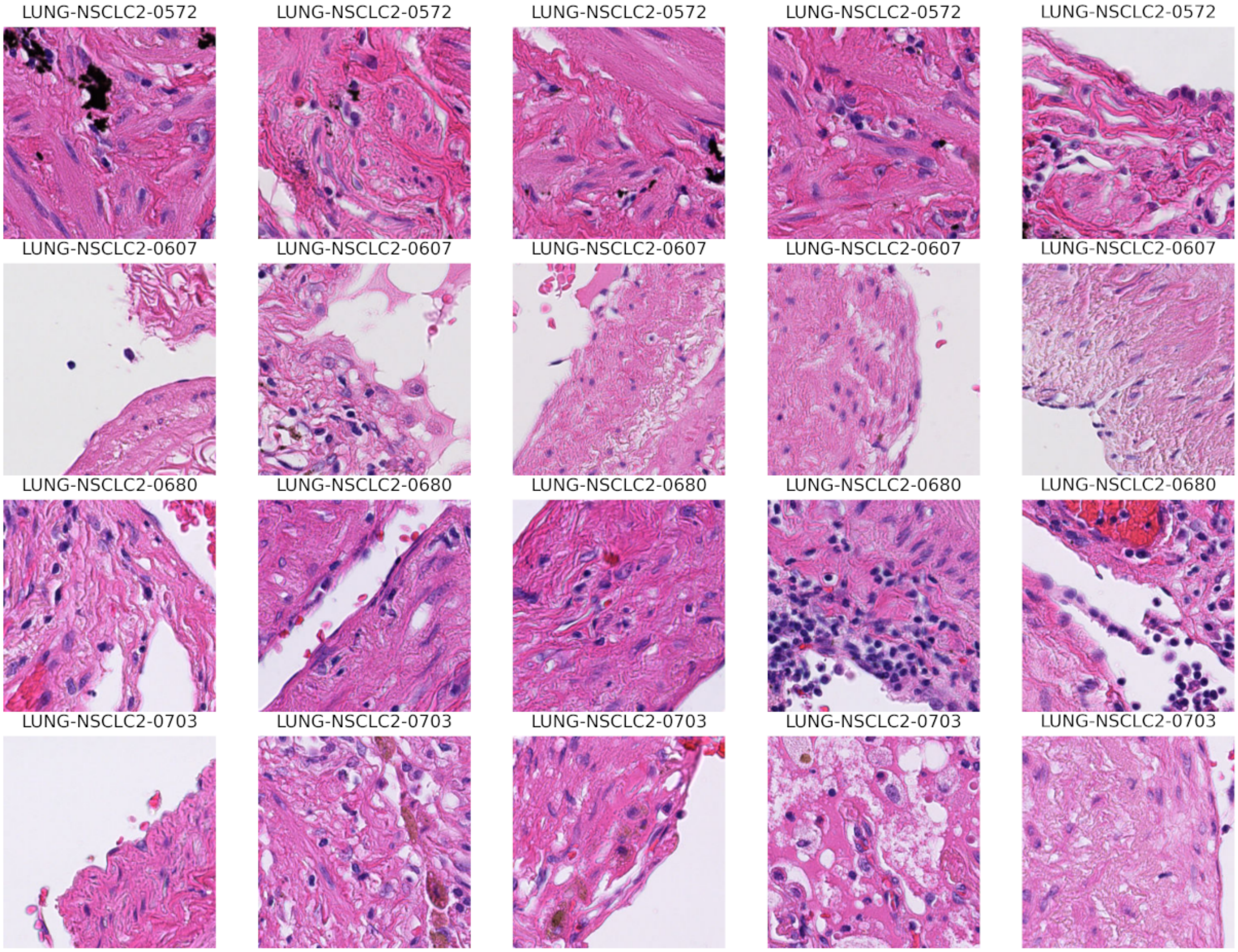


Supplementary Figure 13: Examples of H&E patches from cluster 7 – reactive desmoplastic stroma.

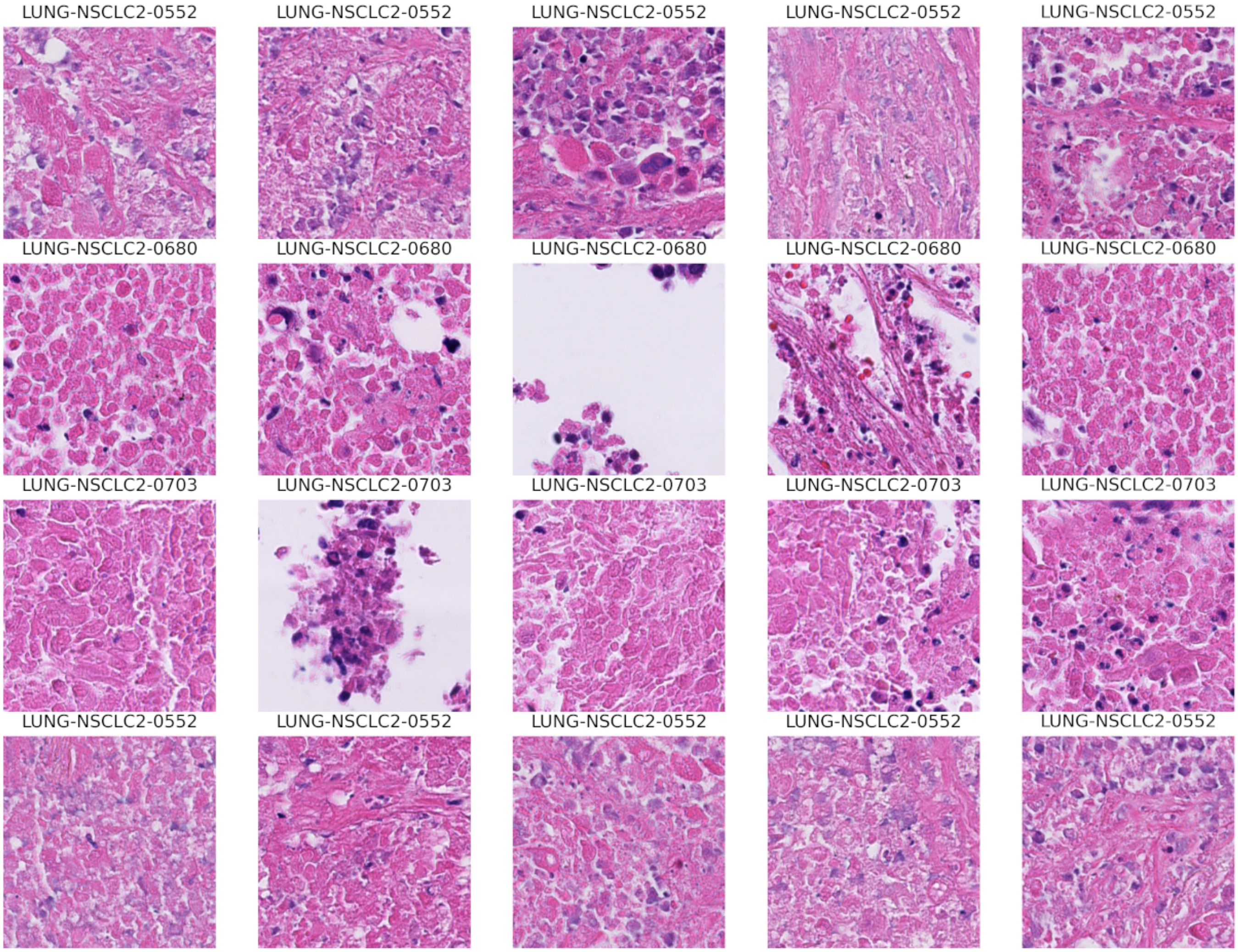


Supplementary Figure 14: Examples of H&E patches from cluster 8 – homogeneous necrotic masses.

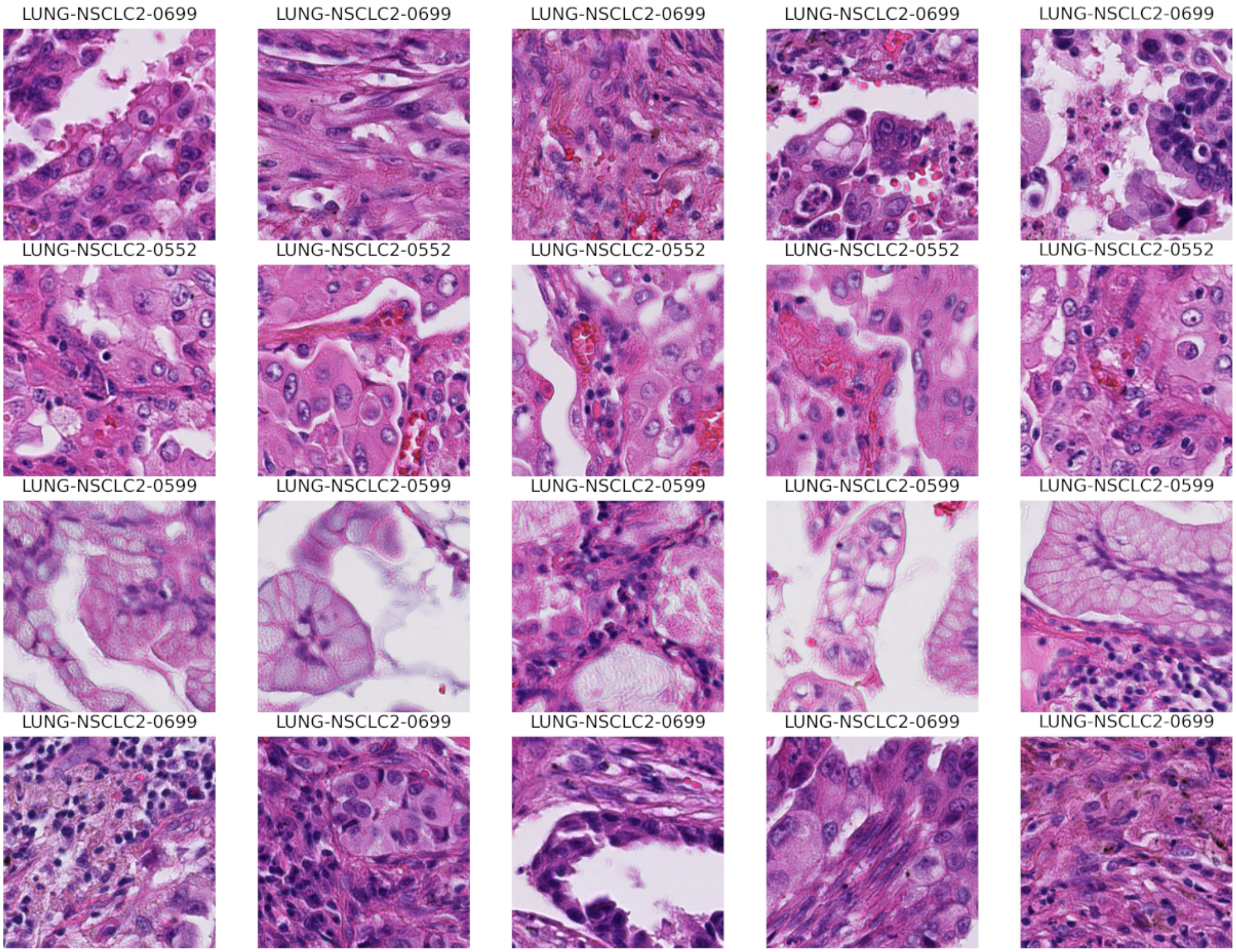


Supplementary Figure 15: Examples of H&E patches from cluster 9 – tumour epithalium-stroma interface.

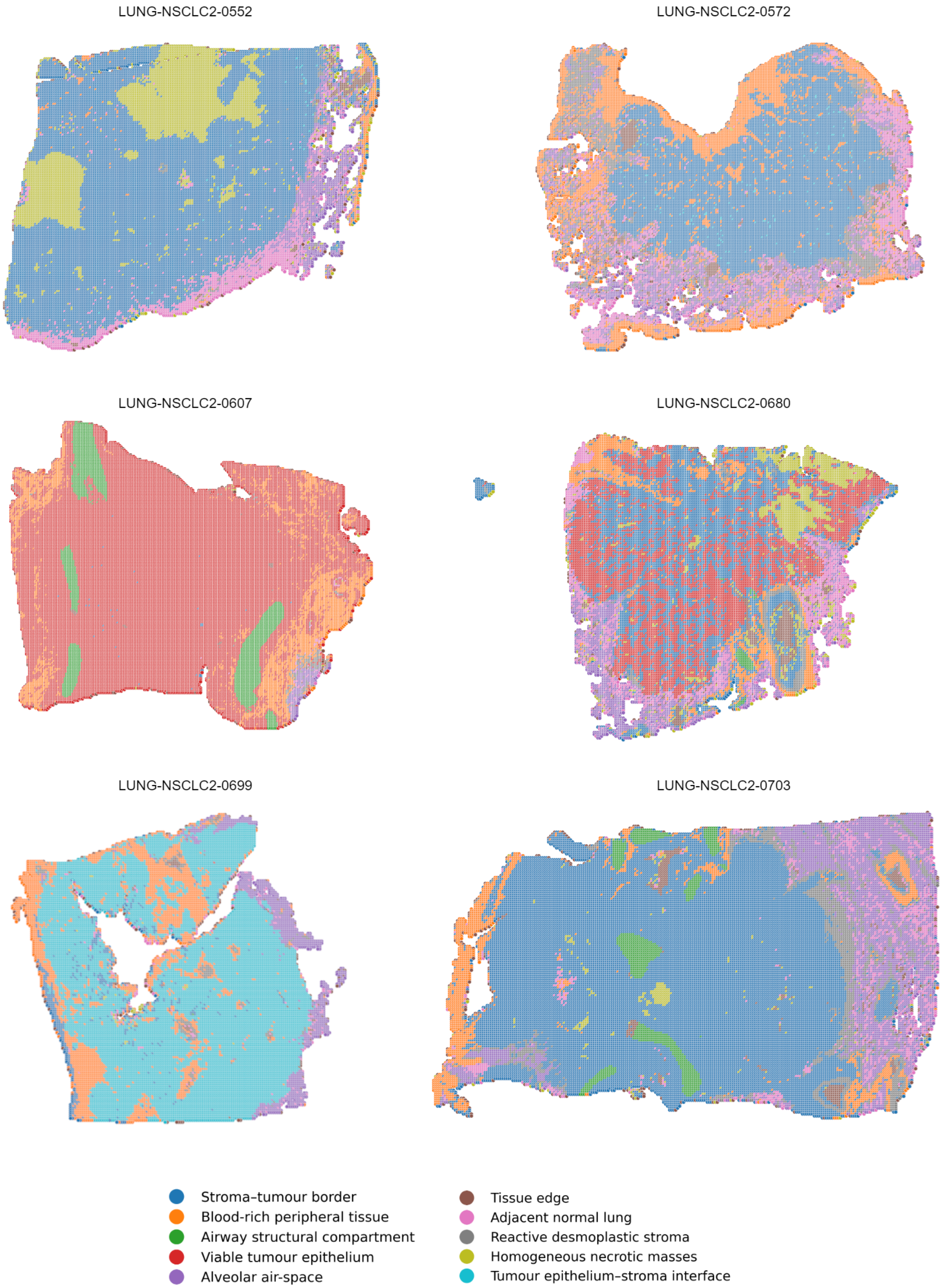


Supplementary Figure 16: Examples of H&E slides coloured by patch assignment to clusters.

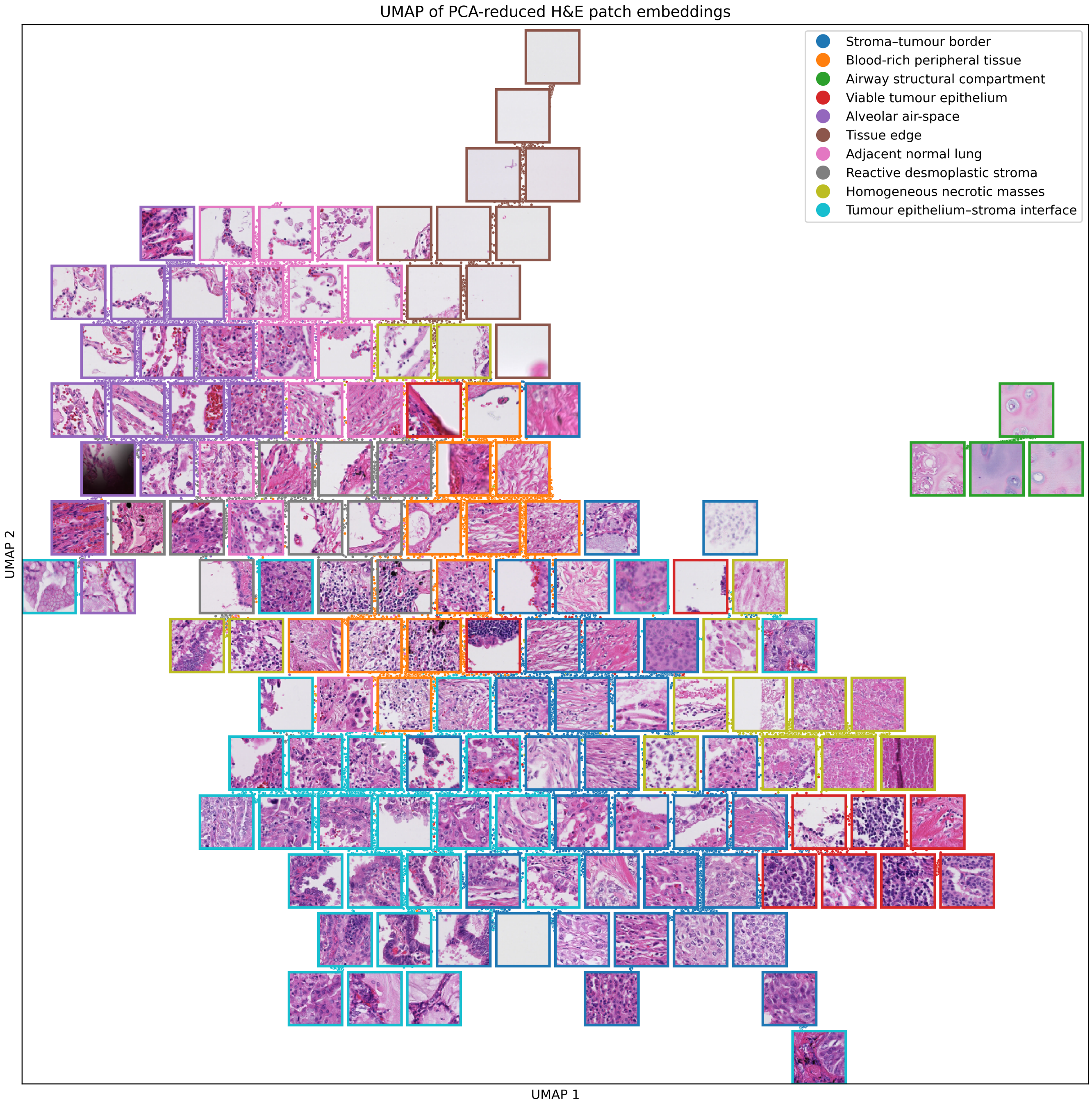


Supplementary Figure 17: UMAP of 40000 sample H&E patches from patients, coloured by their a posteriori assignment in the model, with examples.

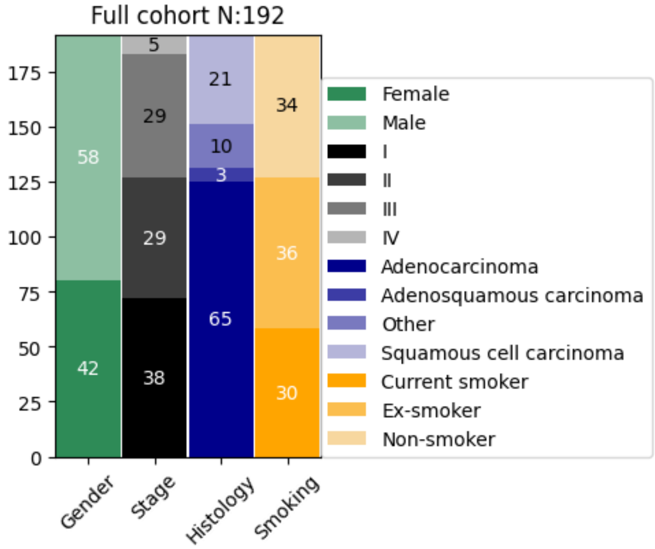


Supplementary Figure 18: summary of the full NSCLC2 cohort by clinical features

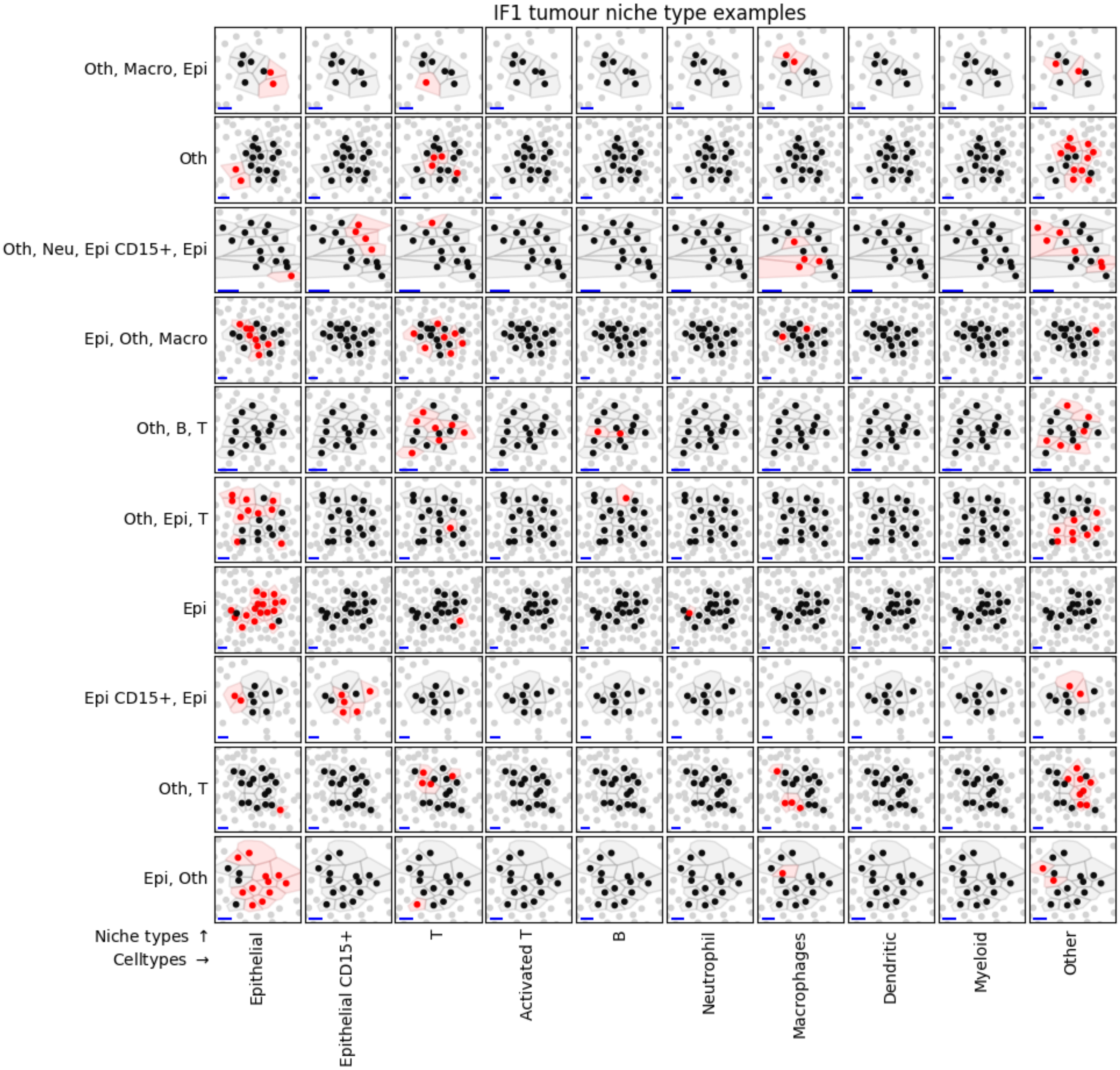


Supplementary Figure 19: Examples of niche types in IF1 tumour regions. Each row corresponds to one niche type, labelled by the celltypes with highest abundances in the niche's profile in decreasing order. One example niche is illustrated per row, with all cells from the niche coloured black, except the cells in red, which have the column's celltype. Blue line represents 10$\mu$m for scale.

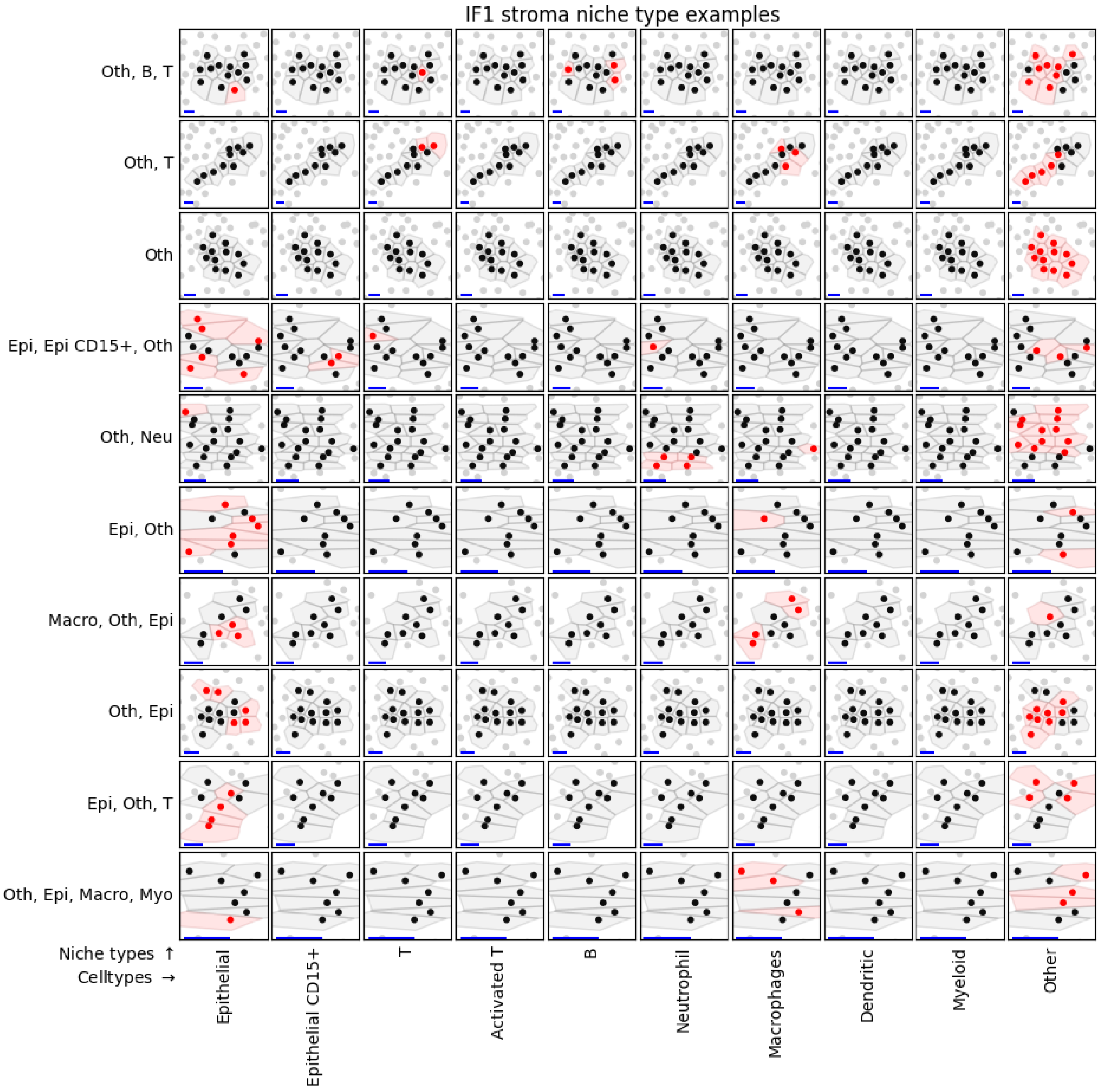


Supplementary Figure 20: Examples of niche types in IF1 stroma regions. Each row corresponds to one niche type, labelled by the celltypes with highest abundances in the niche's profile in decreasing order. One example niche is illustrated per row, with all cells from the niche coloured black, except the cells in red, which have the column's celltype. Blue line represents 10$\mu$m for scale.

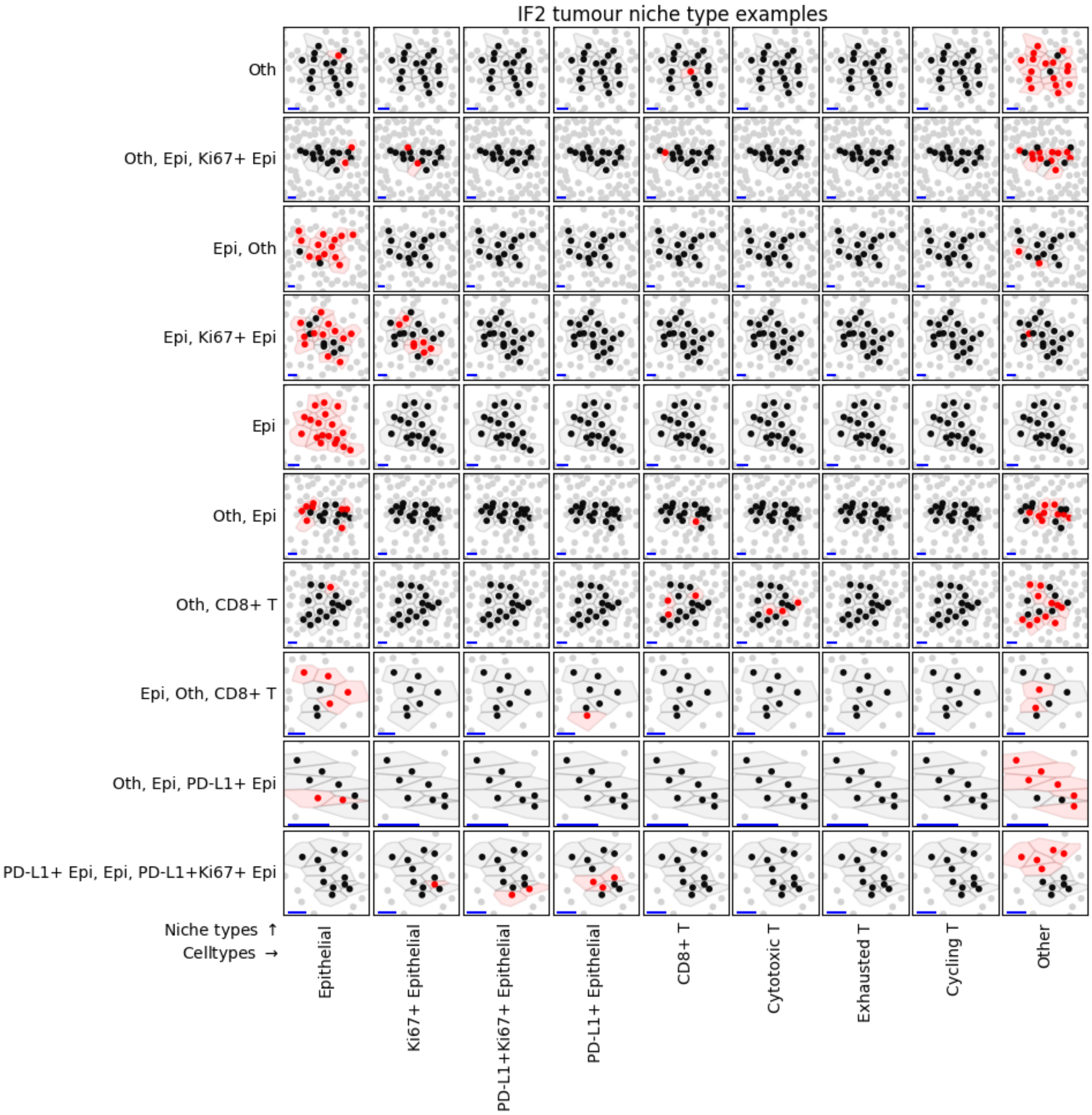


Supplementary Figure 21: Examples of niche types in IF2 tumour regions. Each row corresponds to one niche type, labelled by the celltypes with highest abundances in the niche's profile in decreasing order. One example niche is illustrated per row, with all cells from the niche coloured black, except the cells in red, which have the column's celltype. Blue line represents 10$\mu$m for scale.

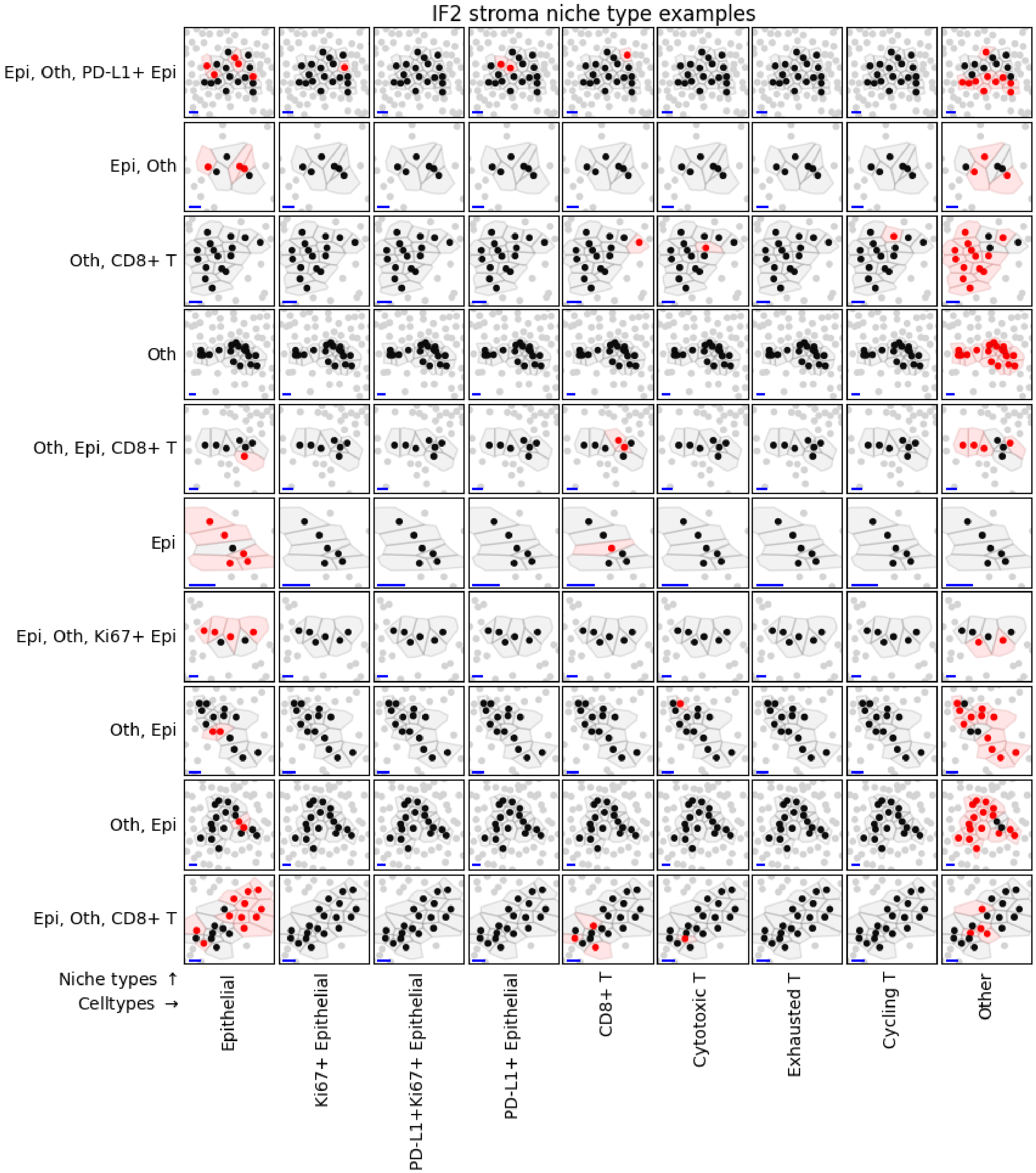


Supplementary Figure 22: Examples of niche types in IF2 stroma regions. Each row corresponds to one niche type, labelled by the celltypes with highest abundances in the niche's profile in decreasing order. One example niche is illustrated per row, with all cells from the niche coloured black, except the cells in red, which have the column's celltype. Blue line represents 10$\mu$m for scale.

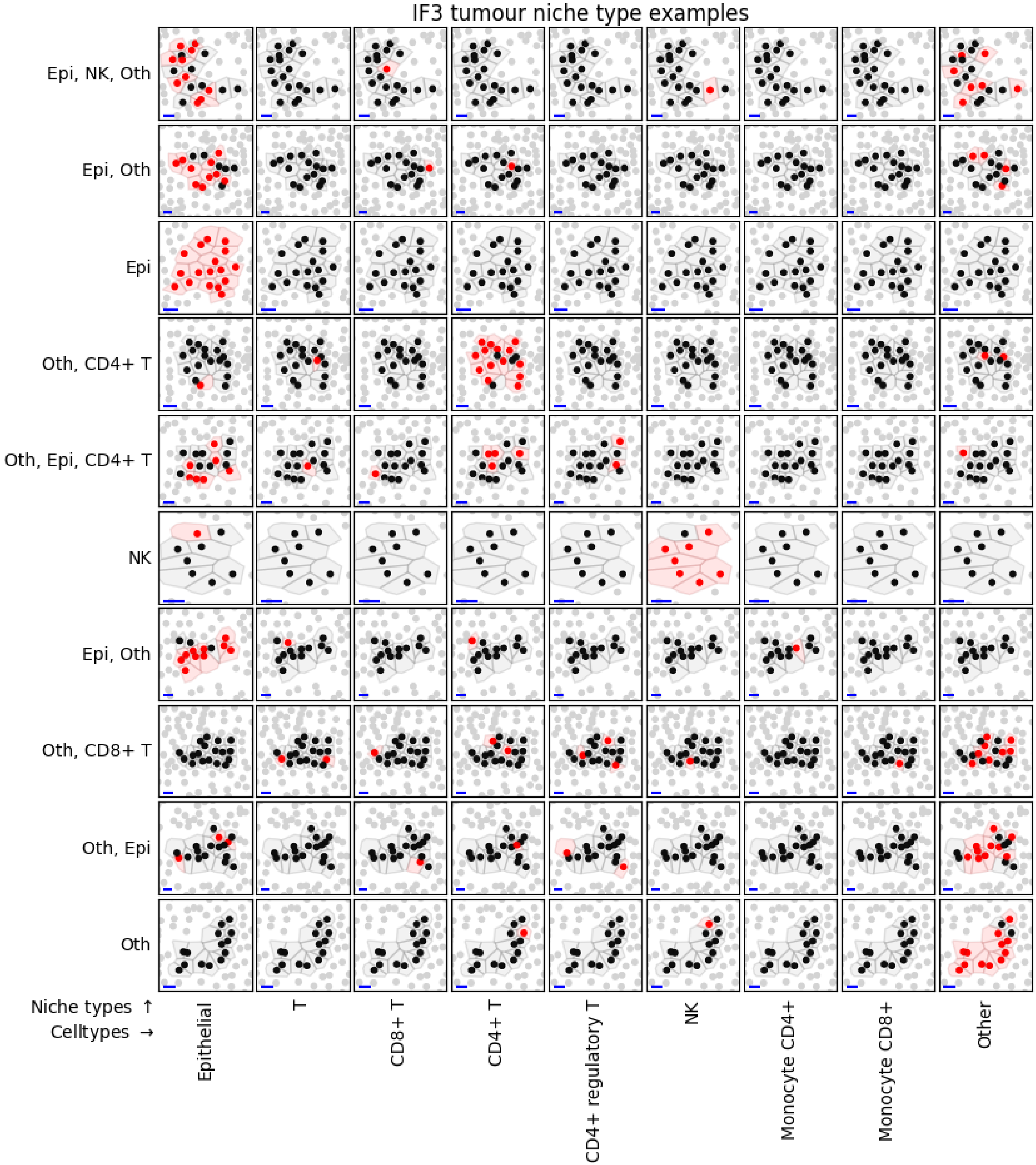


Supplementary Figure 23: Examples of niche types in IF3 tumour regions. Each row corresponds to one niche type, labelled by the celltypes with highest abundances in the niche's profile in decreasing order. One example niche is illustrated per row, with all cells from the niche coloured black, except the cells in red, which have the column's celltype. Blue line represents 10$\mu$m for scale.

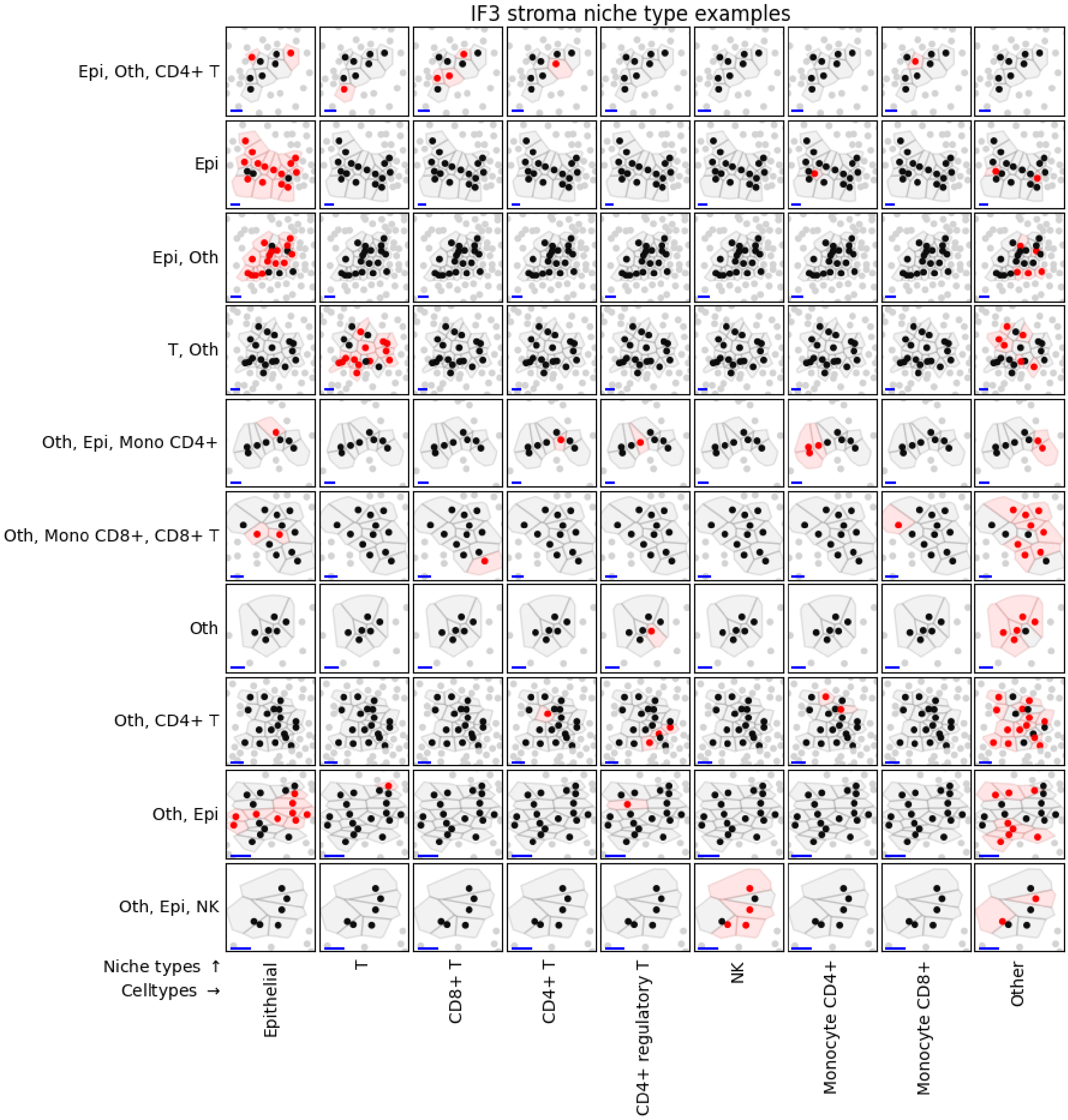


Supplementary Figure 24: Examples of niche types in IF3 stroma regions. Each row corresponds to one niche type, labelled by the celltypes with highest abundances in the niche's profile in decreasing order. One example niche is illustrated per row, with all cells from the niche coloured black, except the cells in red, which have the column's celltype. Blue line represents 10$\mu$m for scale.

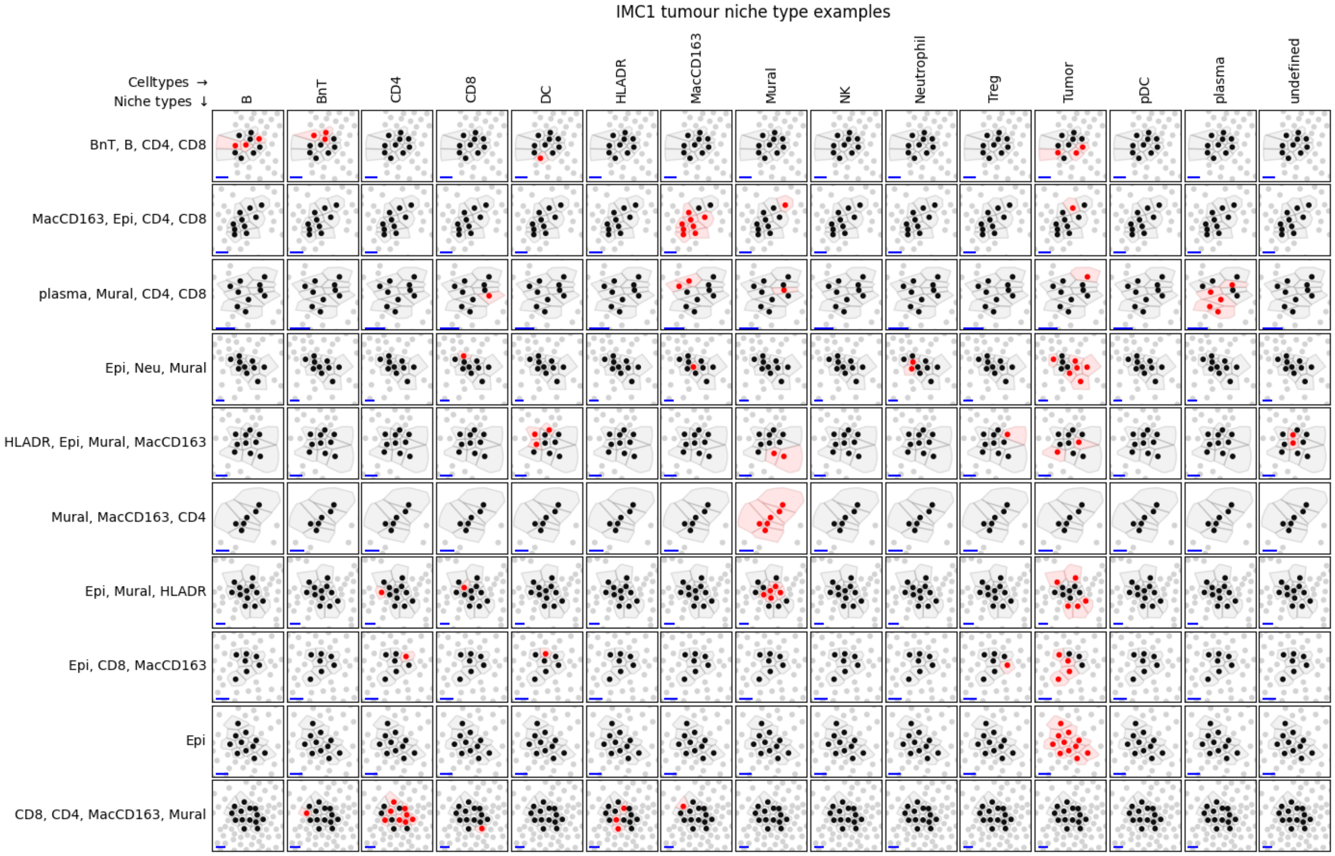


Supplementary Figure 25: Examples of niche types in IMC1 tumour regions. Each row corresponds to one niche type, labelled by the celltypes with highest abundances in the niche's profile in decreasing order. One example niche is illustrated per row, with all cells from the niche coloured black, except the cells in red, which have the column's celltype. Blue line represents 10$\mu$m for scale.

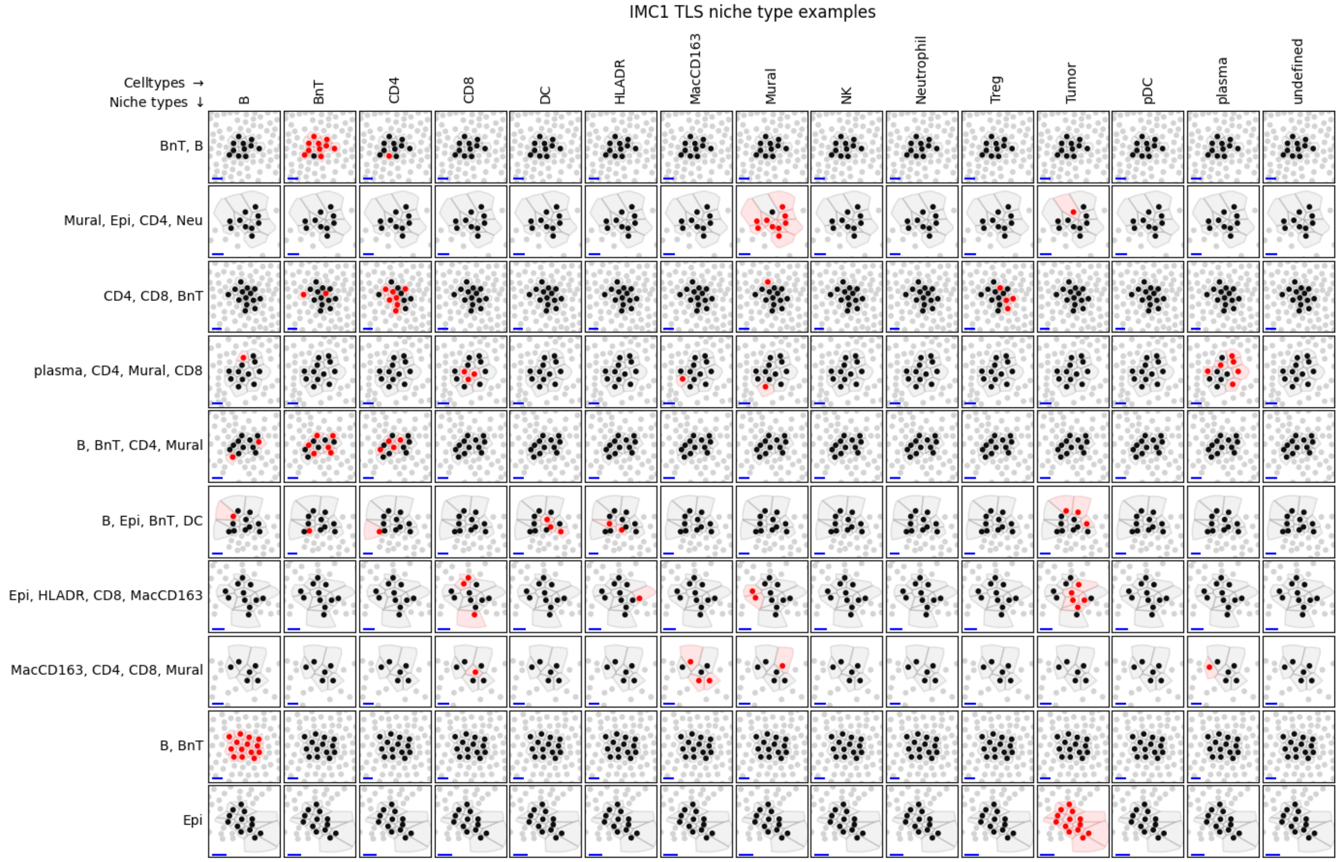


Supplementary Figure 26: Examples of niche types in IMC1 TLS regions. Each row corresponds to one niche type, labelled by the celltypes with highest abundances in the niche's profile in decreasing order. One example niche is illustrated per row, with all cells from the niche coloured black, except the cells in red, which have the column's celltype. Blue line represents 10$\mu$m for scale.

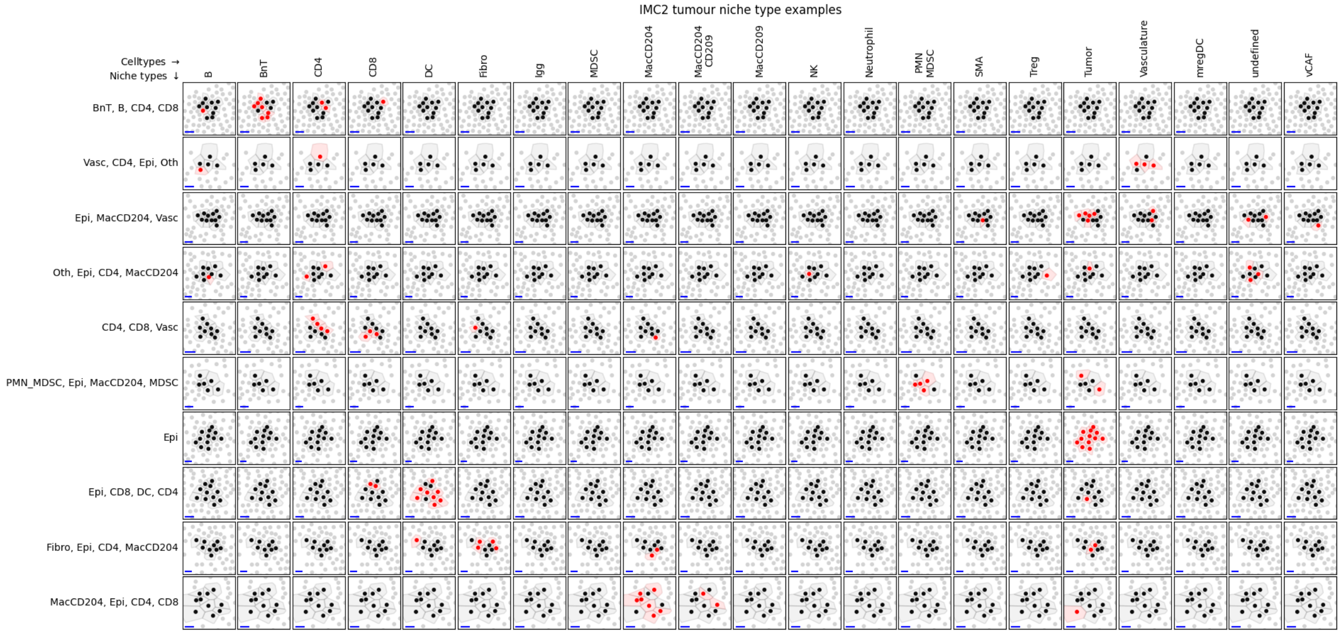


Supplementary Figure 27: Examples of niche types in IMC2 tumour regions. Each row corresponds to one niche type, labelled by the celltypes with highest abundances in the niche's profile in decreasing order. One example niche is illustrated per row, with all cells from the niche coloured black, except the cells in red, which have the column's celltype. Blue line represents 10$\mu$m for scale.
